# Mosaic accumulation of somatic genetic variation and estimates of age in the long-lived reef-building coral *Acropora palmata*

**DOI:** 10.1101/2025.04.11.641527

**Authors:** Trinity Conn, Jessie Renton, Valerie F. Chamberland, Zoe Dellaert, Thorsten B.H. Reusch, Benjamin Werner, Iliana B. Baums

**Affiliations:** Department of Biology, The Pennsylvania State University, University Park, PA; Conservation Research Department, John G. Shedd Aquarium, Chicago, IL, USA; Barts Cancer Institute, Queen Mary, University of London; SECORE International, Miami, 33145, FL; CARMABI Foundation, Piscaderabaai z/n, Willemstad, Curaçao; Department of Freshwater and Marine Ecology, Institute for Biodiversity and Ecosystem Dynamics, University of Amsterdam, Science Park, Amsterdam 1098 XH, The Netherlands; GEOMAR Helmholtz-Centre for Ocean Research Kiel, Wischhofstr. 1-3, 24148 Kiel, Germany; Helmholtz-Institute for Functional Marine Biodiversity at the University of Oldenburg [HIFMB], Im Technologiepark 5, 26129 Oldenburg, Germany; Alfred Wegener Institute, Helmholtz-Centre for Polar and Marine Research [AWI], Am Handelshafen Bremerhaven, Germany; Institute for Chemistry and Biology of the Marine Environment (ICBM), School of Mathematics and Science, Carl von Ossietzky Universität Oldenburg, Ammerländer Heerstraße 114-118, 26129 Oldenburg, Germany

**Keywords:** Somatic Mutations, Corals, Adaptation, Age, Allele Frequency

## Abstract

Somatic genetic variation (SOGV), accumulating during an organism’s lifetime, was traditionally viewed as detrimental rather than adaptive due to links with cancer and senescence. However, in modular organisms like corals, deleterious mutations can be purged at the cellular or polyp level, while adaptive mutations may rise in frequency as polyps create genetically distinct modules. Quantifying the somatic genetic landscape in corals is necessary to understand the role these mutations may have in coral and clonal animal development and evolution. Here, we catalog somatic genetic variation in eight *Acropora palmata* colonies from Curaçao. Whole genomes were sequenced (70-100x depth), documenting mutation variant allele frequency shifts as genets aged. Large numbers of SOGVs were observed in six- to ten-year-old colonies, and inferred mutation rates were used to age a genet of uncertain age to almost a century old. Although mutations were not fixed at the polyp or branch levels, i.e. they always displayed frequencies <0.5 as expected at mutating homozygous sites, their allele frequencies followed a power-law distribution, similar to aging human tissues. No signs of positive selection were found; instead SOGVs in the colony of uncertain age were under purifying selection. In one colony, mutations in 28 samples from along a branch were analyzed using a SNP microarray. Contrary to expectations, genetic and physical distances were unrelated. This observation together with the observed lack of fixation may be explained by a large stem cell population, the de-differentiation or dormancy of stem cells, the contribution of strong purifying selection, or a combination of the previously mentioned. Our findings provide a neutral framework against which to test for module-level selection of genetic variation in corals, explore the relationship between physical and genetic distance within a colony, and apply a somatic genetic clock to colonies of *Acropora palmata.* This work provides necessary fundamental insights into the landscape of somatic mutations in reef-building coral, highlighting the importance of studying these mutations as they may contribute to genetic diversity and adaptability in colonial animals.

## Introduction

Organisms with a modular bauplan are widespread across the tree of life (Reusch, Baums, & Werner, 2021). Clonal species are special modular organisms that consist of a series of repeating modules that can independently reproduce and survive. The modular structure allows for the accumulation of somatic mutations with time without putting the survival of the colony or genet at risk, because death at the module level does not kill the clone or genet. Somatic mutations thus present an understudied source of genetic diversity within modular organisms, and in many modular taxa they can even be inherited across sexual generations including all plants and some basal animals (Vasquez-Kuntz et al., 2022; Antolin & Strobek, 1985). Studies of the generated somatic genetic variation can inform researchers, practitioners, and managers about the life history traits and phenotypic variation in organisms where long lifespans, clonal expansion, and limited genomic resources may have previously prevented such research Hughes & Jackson 1980; Devlin-Durante et al., 2016). Further, such studies help connect demographic processes with evolutionary dynamics in novel ways by expanding our understanding of the role of genetic variation within modules to adaptation and genetic diversity at the population level (Reusch et al., 2021; Schoen & Schultz, 2019; Ranade et al., 2015).

Previously, a stepwise mutation model of neutral somatic mutation accumulation based on microsatellite markers to age genets was first applied in aspens (Ally, Ritland, & Otto, 2010) and then corals (Devlin-Durante et al., 2016). The maximum age detected in the focal species of the study, *A.palmata*, was 838 to 6500 years depending on mutation rate, indicating that coral genets can be as long lived as the oldest known plant genets (de Witte & Stöcklin, 2010).

With advent of next generation sequencing, a novel universal clock for modular organisms was developed based on genome wide SOGV that accumulates and segregates among ramets of a genet through a process of somatic genetic drift (Yu et al. 2020). In modular organisms, a key finding is that fixation among modules can happen in the absence of selection (Yu et al., 2024). This allows for the use of these fixed mutations to determine the age of genets. After an initial lag phase, fixed somatic mutations increase linearly with time, thus behaving in a clock-like manner. The model suggests that the length of the lag phase depends largely on the stem cell population size, the number of founder cells that form new modules and the type and ratio of cell divisions (asymmetric versus symmetric). Therefore, under large stem cell population size (N), fixation will be very slow. Applied to seagrass clones, the model predicted age in extremely old genets of *Zostera marina* (Yu et al., 2024). This model represents an improvement on previous coalescent models, increasing accuracy of age estimates in long-lived clonal organisms. Here, we explore the genome-wide accumulation of somatic genetic variants in modular reef-building corals to describe the prevalence of somatic mutations and their associated variant allele frequencies and test whether this novel somatic clock can be applied to estimate coral genet age.

The somatic genetic clock was developed using mutations identified as fixed from the allele frequency spectra. However, before becoming fixed, most somatic mutations except very few developmental mutations are at low frequencies and must rise in frequency before becoming visible in bulk sequencing. Before fixation as heterozygotes in diploid species, they exist as a wide distribution of frequencies within an organism, characterized as genetic mosaic, as cell lineages with different genotypes co-occur. These variants can either be lost in the population, experience positive selection, or be neutral and shift in allele frequency due to genetic drift, including possible fixation. The likelihood of fixation of neutral variants increases with decreasing size of the stem cell pool that forms new modules. In plants, this stem cell pool is usually small (3-4 cells in Arabidopsis (Burian et al. 2016), 7-12 cells in a seagrass species, Yu et al. 2024) while the stem cell pool size in corals is unknown. A large stem cell pool forming new modules may prevent fixation of somatic mutations even in long-lived genets as it is less susceptible to random somatic genetic drift. In such a case, it is not clear if the somatic genetic clock can still be used to age genets.

As modular organisms grow and accumulate modules over time, one would expect that the accumulation of somatic mutations would increase with spatial distance relative to the founding module. In corals, the situation is more complex as in simple modular organisms such as seagrasses or other rhizomatous clonal plants. Here, these modules are produced at two levels, polyps and colony fragments that can give rise to entire daughter colonies (Highsmith,1982). First, the colony grows through the asexual division and propagation of polyps within a colony. These polyps are the modules relevant to the modeling of the accumulation of somatic mutations – they (usually) can survive on their own when separated from the colony and they are formed by independent divisions, either intratentacularly or extratentacularly (Coates & Jackson, 1987). When embedded in a colony, polyps share resources and signals through a thin layer of tissue (Oren et al., 2001). These polyps can then continue to produce daughter polyps through polyp divisions (Figure 1a), which will accumulate SOGV over time, generating a diverse genetic landscape across a colony of a scleractinian coral. Studies at the polyp level provide insights into the dynamics of allele frequency changes within genets as described above.

**Figure 1:**
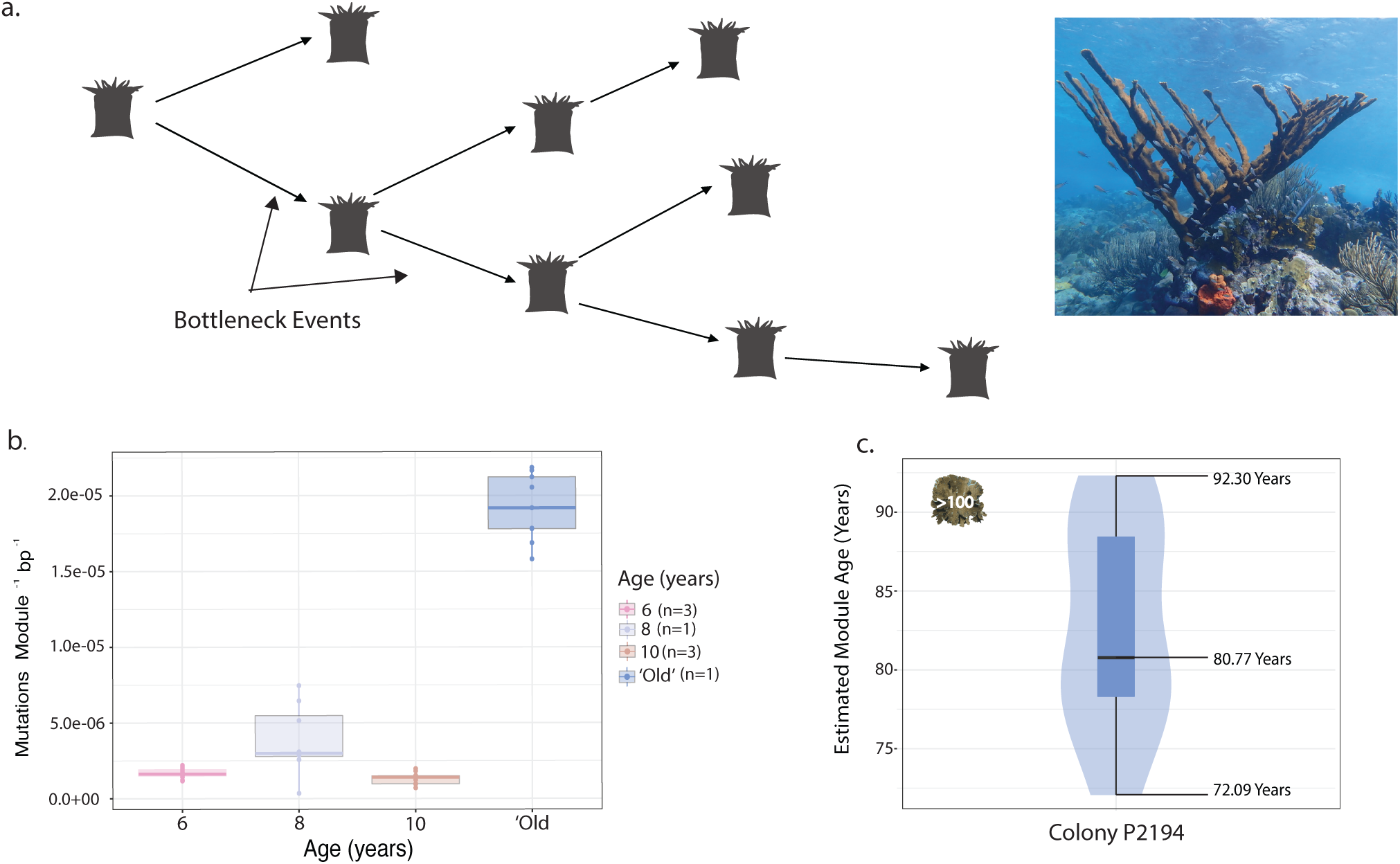
The accumulation of somatic mutations in a colony of *Acropora palmata* to estimate the age of a module. a) The process by which modules within a genet of *A.palmata* can differentiate over the lifetime of the colony. Each time a module (polyp) divides, a subset of cells within a polyp are recruited to contribute to the new module. This results in a bottleneck in the genetic diversity that resulted from mutations within the modules. Overtime, as mutations accumulate in the dividing polyps, modules eventually become genetically distinct, and the pairwise genetic distance between them increases and results in genetically distinct modules, as in a larger colony as pictured. (Photo: Zoe Dellaert). b) Mutation rate per module was calculated for each age class (6, 8, 10 years old and a colony of uncertain age). Each boxplot shows the median (dark-colored line in the central portion of distribution) and the ranges (points) of the mutation rates calculated for all modules within an age class. The top and bottom of the boxes represent the 25% and 75% quartiles of the range of values. c) Distribution of estimated ages based on rate of SOGV occurrence in modules within the ‘old’ colony of uncertain age. The youngest estimated age was 72.09 years old, while the oldest estimated age was 92.30 years old. True genet age likely falls closer to the median of these estimates, approximately 80.77 years old. This is younger than the previously estimated age of this genet estimated using microsatellites, 228-1194 years old (Devlin-Durante et al., 2016)

An ecological extension of this work is to provide much needed accurate estimates of genet age for demographic models. Occasionally, physical disturbance results in the fragmentation of colonies, forming physically independent ramets of a single genet (Baums, Miller, & Hellberg, 2006; Lirman, 2000). Therefore, a small colony may stem from a large old genet or from a recent sexual recruit. Previous attempts at aging have used size, and skeleton cores to date the age of a coral (Tomiak et al., 2016). However, aging the skeleton informs us about the age of the skeleton but not the age of the genet (Irwin et al., 2017). Our difficulties with aging genets also mean that we are lacking crucial information on population dynamics of reef building coral species such as the frequency of sexual recruitment which makes it more challenging to manage these species. Here, we had access to colonies of known age that resulted from coral restoration efforts, a prerequisite to solving the problem of aging coral genets.

In this study, we use deep sequencing of colonies of known age to test the applicability of a previously developed universal somatic genetic clock in corals, quantify the allele frequency spectra of somatic mutations in corals and identify signals of selection or neutrality. Previous work in human cancer and cell biology has identified that in normal cells, the Variant Allele Frequency (VAF) spectra of populations of cells under neutral conditions and constant population size approach a 1/f power law distribution, while those experiencing exponential population expansion follow a 1/f^2^ distribution (Moeller et al., 2024). Moreover, neutrality can be assessed by observing the functional impact of somatic mutations across the genome, where neutral forces should create a pattern of random distribution of mutations across the genome. Deviations from these expectations may indicate positive selection. This paper describes three major aspects of the role of somatic genetic variation in corals: (i) the somatic genetic landscape in *Acropora palmata*, (ii) test the correlation between physical proximity and shared SoGV, and (iii) test the application of a somatic genetic clock through calibration with colonies of known age.

## Materials and Methods

### Sample and Data Collection

Seven *Acropora palmata* colonies of known age were selected from a cohort of outplants monitored by SECORE across two reef sites in Curacao (Sea Aquarium: 12.0839146 ° N, 68.894431° W; Marriott: 12.11888613 ° N, -68.9685206 ° W; Table S1). Colonies represented three distinct age classes: 6 years old (*n*=3 colonies), 8 years old (*n=*1 colony), and 10 years old (*n*= 3 colonies). These colonies are part of cohorts produced under human care during coral spawning in 2011 and 2015. Scientists at SECORE International and CARMABI have monitored these colonies since they were outplanted. These colonies were previously genotyped using the Applied Biosystems Axiom Coral Genotyping Array – 550962 (Thermo Fisher, Santa Clarita, CA, USA) (Kitchen et al., 2020).

Each colony was sampled 5-6 times using coral cutters. Samples consisted of 3-4 polyps each and the location of each sample was noted both with images and in the sampling notes. Samples from the 6-year-old colonies and the 10-year-old colonies were collected in December of 2021, while samples from the 8-year-old colony were collected in September of 2019. Samples (n=9) were also collected from one ‘old’ colony previously estimated to be 228-1924 years old using microsatellite markers (Devlin-Durante et al, 2016). Samples were preserved in 95% ethanol and kept at room temperature until return to Penn State University and then transferred to -20°C. DNA was extracted using the Qiagen DNeasy Kit (Qiagen, MA), following the modified protocol by Kitchen et al., 2020 (dx.doi.org/10.17504/protocols.io.bgjqjumw). All libraries were prepared following the Truseq PCR-Free protocol (Illumina, CA). Samples were sent for paired-end whole genome sequencing with an insert size of 150bp and a target depth of 80x on the NovaSeq 6000 (Table S1).

### Data Processing

Sequencing data was checked for quality using FastQC version 0.11.8 (https://www.bioinformatics.babraham.ac.uk/projects/fastqc/). Sequences were trimmed for quality and adaptor contamination using BBduk version 39.06-1 (https://jgi.doe.gov/data-and-tools/bbtools/bb-tools-user-guide/bbduk-guide/). The following parameters were used to trim the sequences: (trimq=20, minlen=50, maq=20). Sequences were aligned to the *Acropora palmata* genome (Locatelli et al., 2023) using the Burrows Wheeler method using BWA-MEM v0.7.17 with default parameters (Li & Durbin, 2009). Reads were then compressed and sorted using SAMTools version 1.20.0. Duplicated reads were marked and removed using the MarkDuplicates tools in GATK version 4.0.1.2. Properly paired reads with a minimum mapping score of 20 (-q 20) were retained using SAMtools version 1.20.0 (Danecek et al., 2021) Coverage across scaffolds was calculated using bbmap version 39.06-1 (Supplementary Data Table S1).

### Identification of Somatic Mutations

Somatic Mutations were identified using Mutect2 in GATK version 4.0.1.12 using standard settings and the *Acropora palmata* reference genome (Benjamin et al., 2019) using a tumor-normal pairing-based method. All samples within a genet were compared in a pairwise fashion, with each sample treated as the ‘tumor’ sample and the ‘normal’ sample. Identified SNPs (single nucleotide polymorphisms) were then filtered following the standard parameters set out by Mutect2. Mutect2 is able to detect mutations at frequencies as low as 0.01 within the data. As well, only SNPs with a mapping quality of 20 or greater and a minimum base quality of 25 or greater were kept. The total number of SoGVs for each sample within a genet was determined as the sum of all unique mutations discovered for a sample when the sample was classified as the ‘tumor’ in the pairwise comparison. Mutations were filtered using custom scripts in R and only variants identified with a sequencing depth of 60 or higher were retained for future analyses.

### Calculating Substitution Burden

Variant call files (VCFs) containing identified mutations were exported using bcftools version 1.20-0 (Danecek et al., 2021) and loaded into R v.4.3.2. Using a custom R script the mutation load for each sample within a genet was determined as the sum of all unique mutations discovered for a sample when the sample was classified in the ‘tumor’ in the pairwise comparison. Mutations that were found in a ‘tumor’ sample in more than one comparison were removed when estimating substitution burden.

We used the total unique mutation load per module per callable bases in the genome for that module to estimate a mutation rate, as instructed by best practices (Bergeron et al., 2022). Callable bases were calculated as the total bases covered in the output of bbmap. Mutation rate was calculated as:

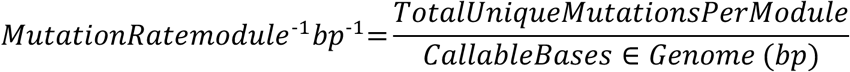

### Estimation of Age

The estimated mutation rate described above was used to calibrate a somatic molecular clock to age the modules of the genet of uncertain age. A mutation rate per module was estimated as the mutation rate year^-1^ in the colonies of known age. The mean mutation rate of occurrence per year was then applied to calculate the age of each module in the genet of uncertain age (Figure 1c.). Age of each module within genet P2194 was calculated as follows:

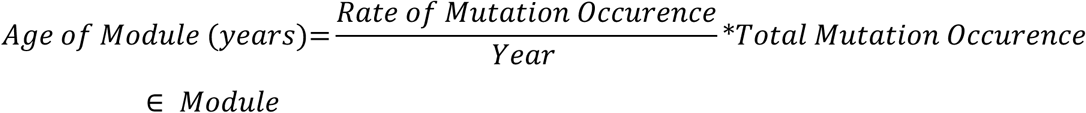

### Distribution of Mutations Across Modules

To investigate how mutations segregate within a colony, the distribution of shared mutations was analyzed. Mutations were classified as ‘private’ if that mutation was not identified in any other sample. Mutations were classified as ‘shared’ if that mutation was identified in greater than one sample for samples treated as tumors. The assumption is that it is more likely that these mutations were shared through shared ancestry rather than the same mutation occurring independently in two tissue samples. A presence/absence matrix of all identified mutations per module was created in R and intersections of mutations were calculated using the R package UpsetR (Conway, Lex, & Gehlenborg, 2017). Mutations were displayed using an UpSet plot that shows the relationship of the mutations across a sampled colony through displaying all intersection sets between modules sampled. Identity and allele frequencies of shared and private mutations were extracted using a custom R script and plotted (Figure 4).

### Co-occurrence Network Analysis

To visualize the relationship between the spatial distribution of genetic diversity (identified somatic mutations) and the physical distance between modules, a co-occurrence network was built for colony M145 (6 years old). A presence/absence matrix was created for all mutations across each sample. A co-occurrence matrix was created by multiplying the matrix against itself transposed in R (co_mat<-t(presabsmatrix)%*% presabsmatrix)). This matrix was then plotted as a network using the R package igraph (Csardi & Nepusz, 2005). Edges indicate the degree of relatedness, where shorter edges indicate that the two modules share more mutations, and longer edges indicate that the two modules that share an edge share fewer mutations between the two of them. The size of nodes in the matrices is indicative of the total number of mutations that module shares with the remainder of the colony. Therefore, the smaller the node, the more distinct the module is from the remainder of the colony. The larger the node size, the more similar the module is to the remainder of the colony (Figure 5b).

### Detecting Population Processes in allele frequencies

Mutation location, reference and alternative bases, and variant allele frequency were exported using bcftools. Using a custom R script, mutations were filtered by depth and by allele frequency. All calls per genet were consolidated into one dataset. Mutations were then classified as ’fixed’ or ’mosaic.’ Fixed mutations were defined as mutations identified with a minimum variant allele frequency of 0.5, while the reference allele had a variant allele frequency less than 0.01 (Yu et al., 2024). Any mutations with a variant allele frequency between 0.01 and 0.49 within the ‘mutant’ individual were categorized as mosaic mutations.

Studies of healthy human somatic cell populations suggest that allele frequencies of cells under neutral processes should demonstrate a relationship of 1/f or 1/f ^2^ in a clonal population undergoing a stable phase (1/f) or a growing phase (1/f^2^) under neutral conditions (Moeller et al., 2024). Given the lack of fixed mutations, even in a colony of theoretically high age, the data was analyzed assuming it follows a model similar to healthy tissues in humans. Therefore, under neutral conditions it is predicted that the allele frequencies of mutations identified in the coral tissue should follow the 1/f^2^ power law distribution (Moeller et al., 2024, Williams et al., 2016). To test the deviation of the allele frequency distributions from a neutral distribution, the R package neutralitytestR (Williams et al., 2016) was used to assess whether the allele frequencies of the genets deviated significantly from the expected neutral distribution. Significance of deviations are assessed by three statistics: the Kolmogorov-Smirnov statistic, the area under the curve (AUC), and a linear regression between the cumulative distribution functions (CDF) of the empirical data and a simulated neutral distribution.

We used the R package MOBSTER (Caravagna et al., 2020) to further deconvolute the clonal clusters present within the genets analyzed to detect whether there were any signals of individual sub clonal clusters. The R package MOBSTER was designed to identify patterns of clonal expansion in bulk sequencing of cancer cells. MOBSTER can identify signals of positive selection as well as capture the neutral evolutionary forces that shape the allele frequency spectra within cancer cell populations. MOBSTER uses a finite mixture model with mixed distributions to identify clonal expansions and neutral dynamics with a 1/f^2^ tail. The model then determines the optimal k to suggest if a sample is monoclonal or polyclonal.

### Annotating Identified Variants

To annotate any functional impact of somatic mutations discovered, the R package dndscv (Martincorena et al., 2017) was used. The functional impact of all mutations located in coding regions was established as either a synonymous mutation, a nonsense mutation, or a missense mutation. Under neutrality, the proportion of mutations identified in coding regions should be statistically like the proportion of the entire genome that is made up of coding sequences. A chi-square test was performed to determine if the observed proportion of mutations discovered in coding sequences was less or more than expected under neutral conditions. All statistical analyses were performed using R v.4.3.2.

### Analyzing Spatial Distribution of Somatic Mutations using a SNP array

#### Sample and Data Collection

In 2019, one large branch of a colony in Curacao was sampled (Blue Bay, 12.13492 , -68.9871) (Table S2). A transect was placed along the branch, and samples were taken every 10 cm along the transect. Samples were also taken at the tips at the end of the branch (Figure 5a & Figure 5.b). These samples were preserved in 95% Ethanol for processing on the StagDB array. DNA was extracted using the Qiagen DNeasy Kit (Qiagen, MA) (**dx.doi.org/10.17504/protocols.io.bgjqjumw**) and sent to ThermoFisher for assaying on the Applied Biosystems Axiom Coral Genotyping Array – 550962 (Thermo Fisher, Santa Clarita, CA, USA).

#### Data Processing

Raw data from the SNP array were analyzed using the Axiom ‘Best Practices Workflow’ using the default settings. Settings were set so that the sample Dish QC> 0.82, plate QC call rate > 97, SNP call-rate cutoff > 97, and the percentage of passing samples > 95. Resulting genotyping files were converted to variant caller format (VCF) using the bcftools plugin affy2vcf (https://github.com/freeseek/gtc2vcf). The VCF was loaded into R v.4.3.2 for downstream analyses.

#### Identification of Somatic Mutations using a SNP array

All probes with missing data were removed from analysis. In this dataset, somatic mutations were assumed to be the minor allele within a genet sampled. Loci with somatic mutations were identified as any locus where not all samples within the genet carried the same allele call at that site. The mutated allele was the minor allele within the group of samples, while the major allele was assumed to be the ancestral allele. This is consistent with previous assays of somatic mutations in *Acropora palmata* using a SNP array (Vasquez Kuntz et al., 2022).

#### Genetic and Distance Correlation

To test if the accumulation of somatic mutations followed expectations that genetic and physical distance should correlate, two methods were used. First, based on the images taken during sampling, distance between samples along the transect was determined. Prevosti’s genetic distance was calculated between all samples using the R package poppr (Kamvar, Tabima, & Grünwald, 2014), derived from the set of probes identified to contain mutations. A linear regression was run using R v4.3.2 between genetic and physical distance.

## Results

### Calibrated substitution burden to estimate the Age of a genet of Acropora palmata

To first understand the overall contribution of somatic mutations to the genetic diversity of *Acropora palmata*, the overall rate of SoGV accumulation was calculated. Overall SoGV accumulation was calculated as the rate of occurrence of mutations per module per base pair (Table 1). The mean substitution rate of detected mutations across all aged colonies equaled 2.798 e^-7^± ± 1.977e^-7^ mutations bp^-1^ module^-1^ year^-1^.

**Table 1:** Average substitution Rate Across Age Classes.

| Age | Average Substitution Rate (mutations $\text{bp}^{-1} \text{ module}^{-1}$) | ± SE (mutations $\text{bp}^{-1} \text{ module}^{-1}$) |
| --- | --- | --- |
| Six | $1.687 \times 10^{-6}$ | $3.227 \times 10^{-7}$ |
| Eight | $3.854 \times 10^{-6}$ | $2.327 \times 10^{-6}$ |
| Ten | $1.313 \times 10^{-6}$ | $3.900 \times 10^{-7}$ |
| Uncertain (>100) | $1.919 \times 10^{-5}$ | $2.225 \times 10^{-6}$ |

Using the mean substitution burden across colonies of known age, the age of the modules across the old colony of uncertain age was estimated and ranged from 72.09 - 92.30 years (Figure.1c.), markedly lower than previous estimates using microsatellites of 228-1924 years (Devlin-Durante et al., 2016).

### Distribution of Variant Allele Frequencies

Under the given filtering conditions, none of the SoGV was fixed. Fixed SoGV were defined as those with a read depth of at least 60x, a variant allele frequency of at least 0.50, and a normal (reference) allele frequency of 0.01. Although previous work has established the use of the variance around the fixed allele frequency of 0.5 (Yu et al., 2024), given no mutations were found at a frequency higher than 0.5, this method was deemed inappropriate in this study.

Next, allele frequency distributions within genets were deconvoluted using MOBSTER to study the effects of drift and identify any putative subclones that have expanded within genets due to selection. If allele frequency distributions were mostly shaped by drift, we expected a higher frequency cluster of mutations that became fixed over time and a low frequency tail of recent mutations. Under selection, a third cluster at mid-frequency would be expected containing alleles with a fitness advantage. The best model as determined by Integrated classification likelihood (ICL) and/or Akaike information criterion (AIC) detected two clusters present in all genets. These clusters represent the low frequency mutations, and mutations that have risen to higher frequency. The mean of cluster 1, higher frequency variants, was highest in the old colony (0.25). (Figure 3). This analysis confirms the lack of a third cluster which would indicate the presence of a subclone under selection.

The variant allele spectra of the colonies within the three classes of known age (six, eight, ten years old) did not diverge significantly from the expected neutral 1/f distribution (AUC =0.0273;0.0029;0.0584, K-S Distance=0.0895;0.0751;0.1314, Mean Distance between CDFs= 0.0519;0.0337;0.0524, R^2^ = 0.9790;0.9886;0.9872). (Figure 3). In contrast, the variant allele spectra of the modules within the ‘old’ colony did significantly diverge from the neutral 1/f^2^ distribution (AUC=0.1423, K-S Distance=0.2671, Mean Distance between CDFs=0.1144, R^2^ = 0.9581). Here, the distribution was closer to 1/f^2^ at higher allele frequencies (Figure 3), possibly evident of other demographic processes or selection with age, although more evidence is needed to make further conclusions.

While there was no fixation of somatic mutations across all colonies analyzed, the density of mutations found in the mosaic region (allele frequency of 0.1 to 0.5) increased with age (Figure 2). Comparisons of the number of mutations in the mosaic region across all age classes were significantly different (p<0.01) (Figure 2b.) and the calculated K-S distance value increased with age (Figure 2b) in a non-parametric two-way Kolmogorov Smirnov test. Further, with increasing age, the greatest density of mutations was found at higher allele frequencies significantly increased (Kolmogorov-Smirnov, p<0.05) (Figure 2).

**Figure 2:**
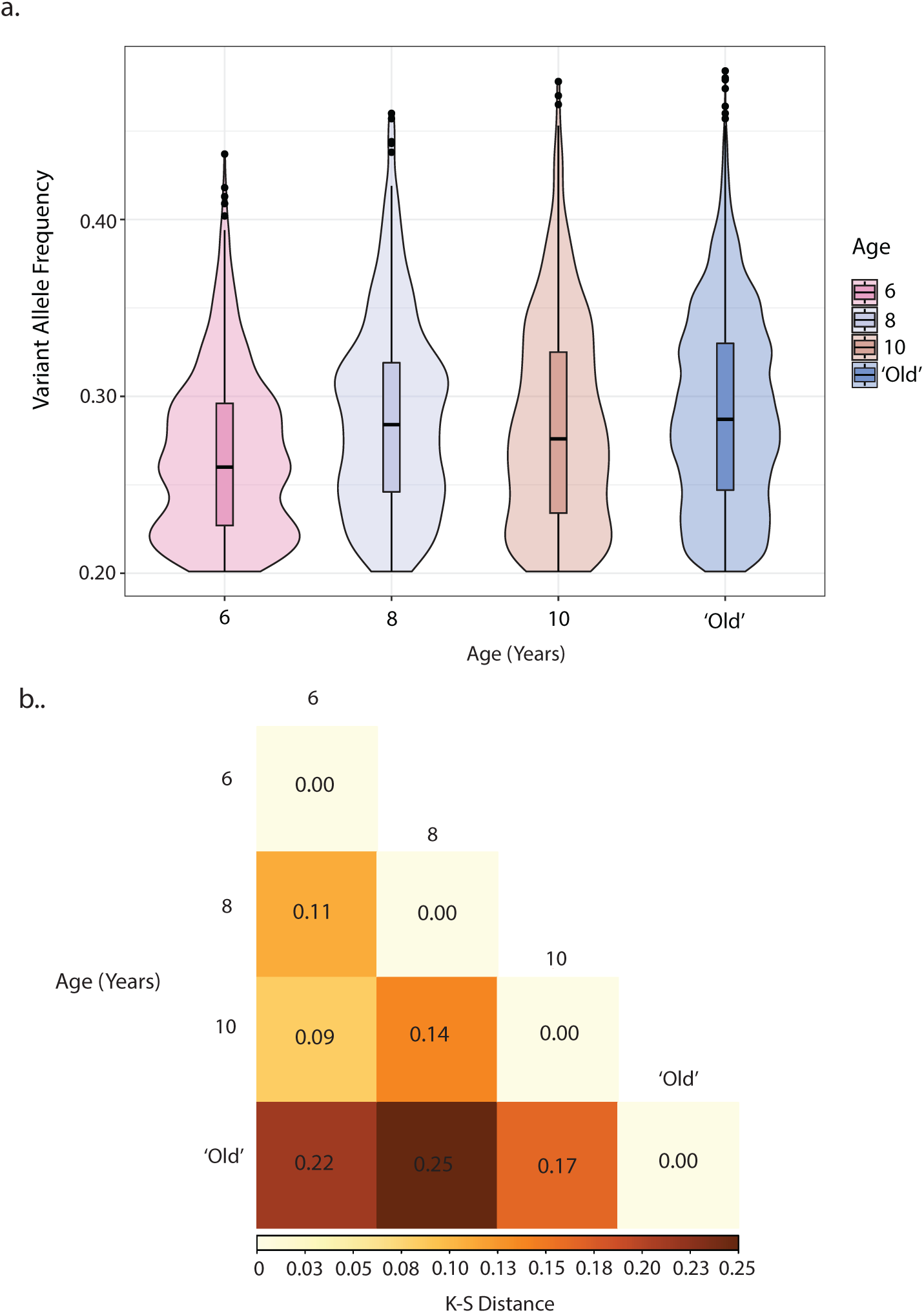
Shifts in allele frequency with age in *Acropora palmata*. (a) Violin plot of variant allele frequencies of mosaic mutations in *A.palmata* with age. Density of mutations at higher frequency increased with age, with the highest density of mutations at high frequency in the ‘old’ colony of uncertain age (blue). Violin plot of allele frequencies in the mosaic state. Only allele frequencies between 0.2 and 0.5 are shown. The boxplots within the violin plots show the median, the range of allele frequencies, and the 25% and 75 % quartiles. Width of violin plot emphasizes distribution of allele frequencies within these quartiles. (b) Kolmogorov-Smirnov distances of all pairwise comparisons between allele frequency distributions from different age classes shown in a). All comparisons were significantly different (p<0.05). Darker colored squares indicate a greater K-S distance. The Kolmogorov-Smirnov test statistic is the maximum distance between the cumulative distribution functions of the two distributions being compared.

**Figure 3:**
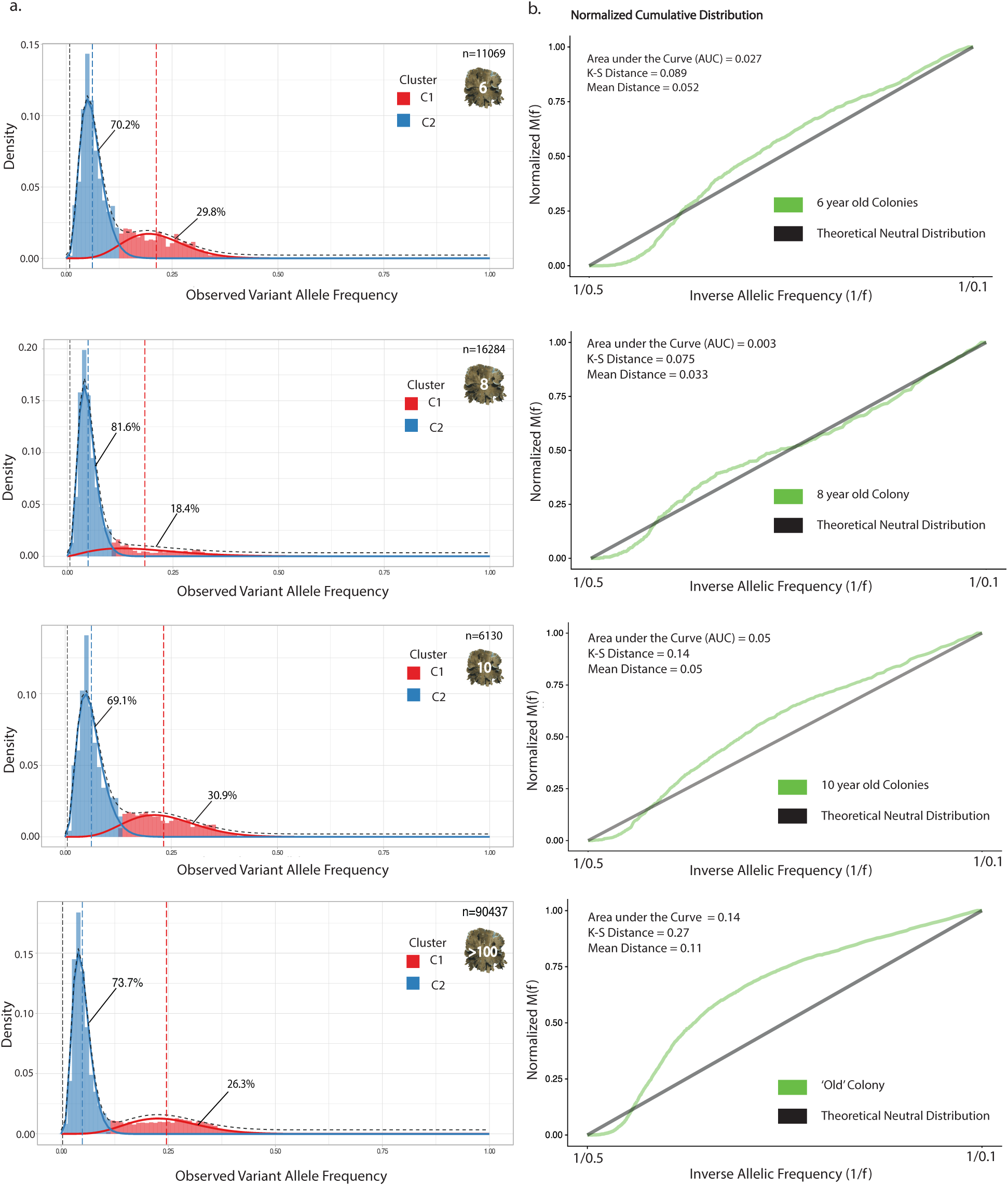
Population dynamics of somatic mutations in *A. palmata*. No clear signal of positive selection was detected across all four colony age classes analyzed in this study. (a) Cluster deconvolution of allele frequency spectra across all four colony age classes. MOBSTER detected two subclones per colony regardless of age. Each colony was made up of mostly polyclonal at low allele frequencies, with one subclone arising to higher allele frequencies. Cluster 1 within each age class is in red, and cluster 2 within each age class is in blue. Dotted line of each cluster’s color represents the mean variant allele frequency of that cluster. (b) Identifying signals of neutrality in variant allele spectra. Each plot represents an age class. Black line is the normalized cumulative distribution function of the theoretical 1/f neutral distribution. Green line within each plot is the normalized cumulative distribution function of the empirical allele frequency within each age class. Statistics of significance for each age class are listed within each plot for each age class.

### Homogeneity of genetic diversity within a colony of Acropora palmata

The majority of identified mutations were private alleles. Mutations were shared among multiple modules in all colonies except for colony S15 (Table S1). Allele frequencies across intersection sets shifted towards higher frequency mutations with more intersection sets (Figure 4c). A co-occurrence network of mutations in colony M145 (Figure 4b.) confirms this analysis, indicating that some modules are more homogeneous with the rest of the population than other modules.

**Figure 4:**
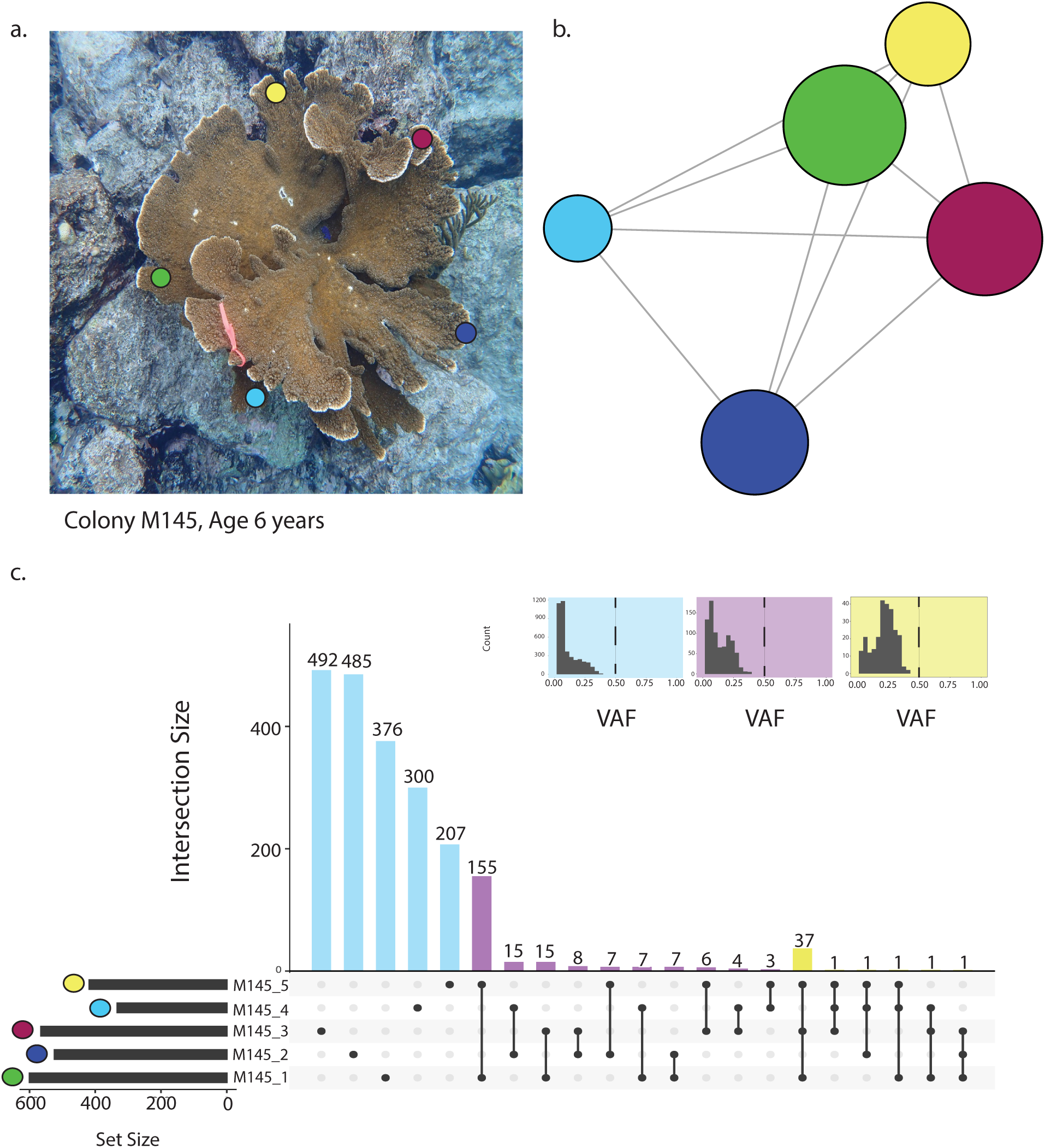
Homogeneity of genetic diversity within a genet of *Acropora palmata.* **(a)** Colony M145, a 6 year old colony, was selected to display the relationship between the physical distribution of modules within a genet and the distribution of mutations across modules. Photo was taken by Zoe Dellaert. Colony M145 is approximately 60 cm in diameter. Colored circles on the colony image (green, yellow, maroon, light blue, dark blue) indicate sampling locations on the colony, and the location of modules analyzed in this study. (b) A co-occurrence network of the mutations displays the connectivity of stem cells within genet M145. The nodes represent the modules, colored by their sampling location as in a) . Larger nodes indicate higher connectivity with other modules. Edge length is correlated with the number of shared mutations, longer edge lengths indicating fewer mutations are shared between the two modules. Module M145_1 (green) has the highest connectivity with other modules, while the light blue module had the least connectivity with other modules, indicating a more unique evolutionary history. (c) UpSet plot of intersection sets of mutations across all modules within colony M145. Modules are labeled by color as they were in a). Intersection sets were categorized as private mutations (light blue bars), mutations shared between two modules (purple bars) and mutations shared with 3+ modules (yellow bars). Allele frequencies of mutations within each category of intersection set were extracted and plotted in the upper right hand of the plot. The background of each plot of observed allele frequencies is color coded by mutation category. A black dotted line represents a variant allele frequency of 0.5.

### Allele Frequency Distributions

The distribution of allele frequencies of private mutations encompasses primarily low frequency variants (Figure 4c) (Supplementary Data Table S3). The next largest group of mutations were mutations shared by two modules (intersection sets). The distribution of allele frequencies of mutations shared between two modules contained both mosaic mutations and low frequency mutations (Supplementary Data Table S3). The distribution of allele frequencies of mutations that were shared between three or four modules were primarily low frequency and mosaic mutations (Supplementary Data Table S3).

We expected that genetically distinct subclones arising during the growth of the colony should be geographically constrained to near the module in which they originated. This expectation was not met in any of the colonies analyzed in this study. The majority (>85%) of identified mutations across all genets were private alleles, only detected in one sample in a colony. Mutations that were shared and retained across regions of the colony are part of higher frequency subclones maintained and propagated across the genet and were also likely generated earlier in the history of the genet, in a shared common cellular ancestor (CCA) of multiple modules . There is an overall shift to higher allele frequencies as mutations are shared with a higher number of modules. However, even low frequency mutations are represented within all intersection sets. Since it is unlikely that the same mutations have arisen in multiple modules independently, some of these mutations are being maintained at low frequency, even after propagation to multiple modules within a genet.

Further evidence for sharing of mutations across a colony of *Acropora palmata* regardless of their geographic origin emerged from an analysis of somatic mutations using a different method, a 19,696 SNP microarray. There was no significant linear relationship between genetic and physical distance (R^2^= 0.0714) along the branch of the colony assayed using the SNP array (Figure 5). There was also discordance between the neighbor joining tree produced by the pairwise genetic distances and the neighbor joining tree produced by the pairwise physical distances. The path-score difference between these two trees was 19.570.

**Figure 5:**
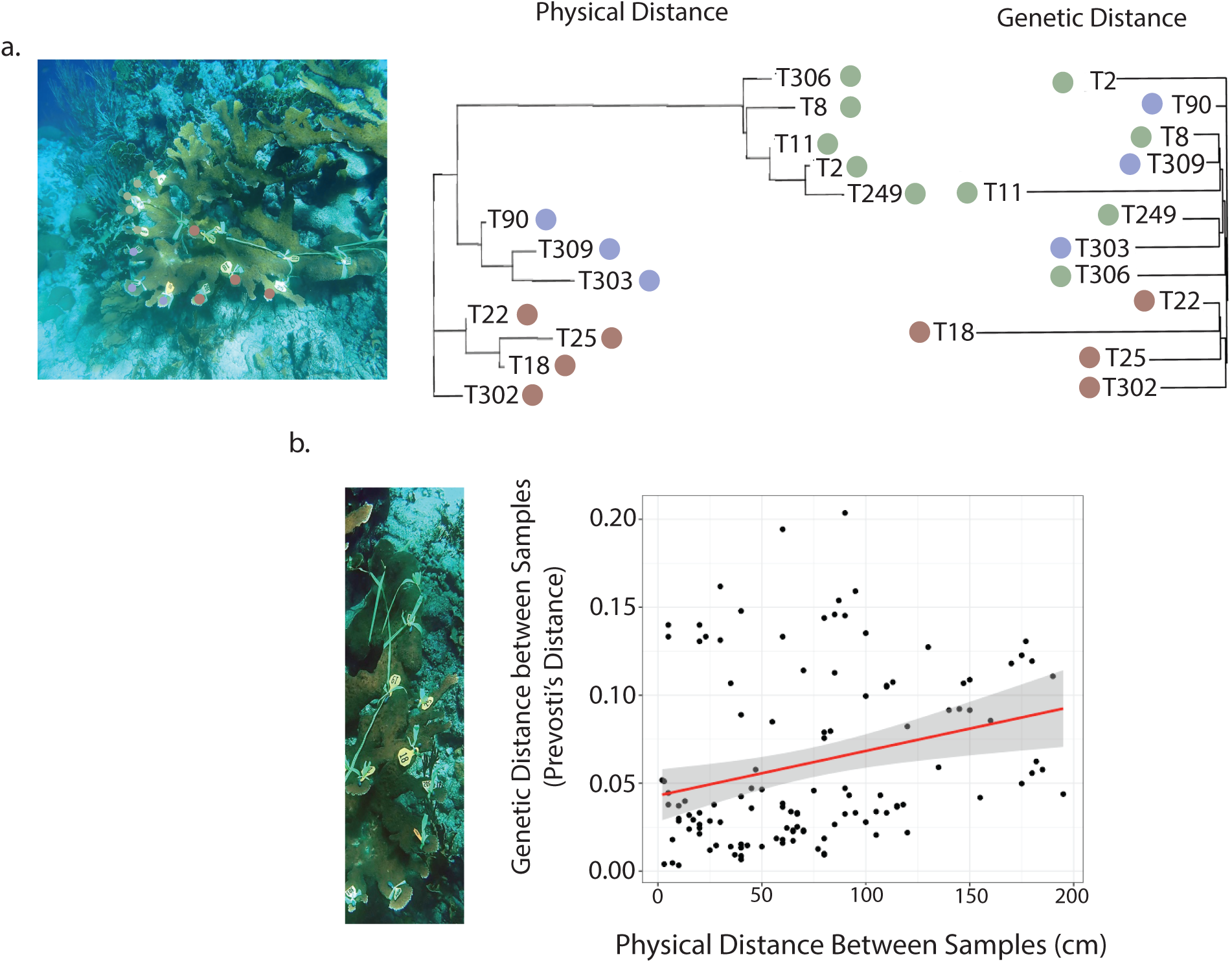
Genetic distance between modules does not correlate with their physical distance along a branch of A. palmata. **(a)** ends of a branch of *Acropora palmata* show discordance between the physical distance (in cm) and the genetic distance (Prevosti’s genetic distance). A neighbor joining tree built using the pairwise physical distance between the tips of the branch. Green, blue and red dots indicate major physical branching events within the colony, creating clusters of closely related growing edges. The pairwise Prevosti’s genetic distances were used to generate a neighbor joining tree of the tips sampled. Physical distance and genetic distance generate different topologies. (b) Linear regression between the Prevosti’s genetic distance and the physical distance in cm along the transect on a large branch of an old colony. There was weak association between the physical distance and the Genetic distance between samples (R^2^= 0.072, p=0.0031).

### Functional Consequences of Somatic Mutation Accumulation

For all samples with at least one mutation detected in coding regions, there was significant depletion of mutations in coding regions (χ^2^, p<0.05) (Figure 6). The proportion of mutations discovered in coding regions across all age classes was never greater than 0.0267 (Data Table S5), significantly reduced relative to the total proportion of coding regions in the entire genome (0.1300).

**Figure 6:**
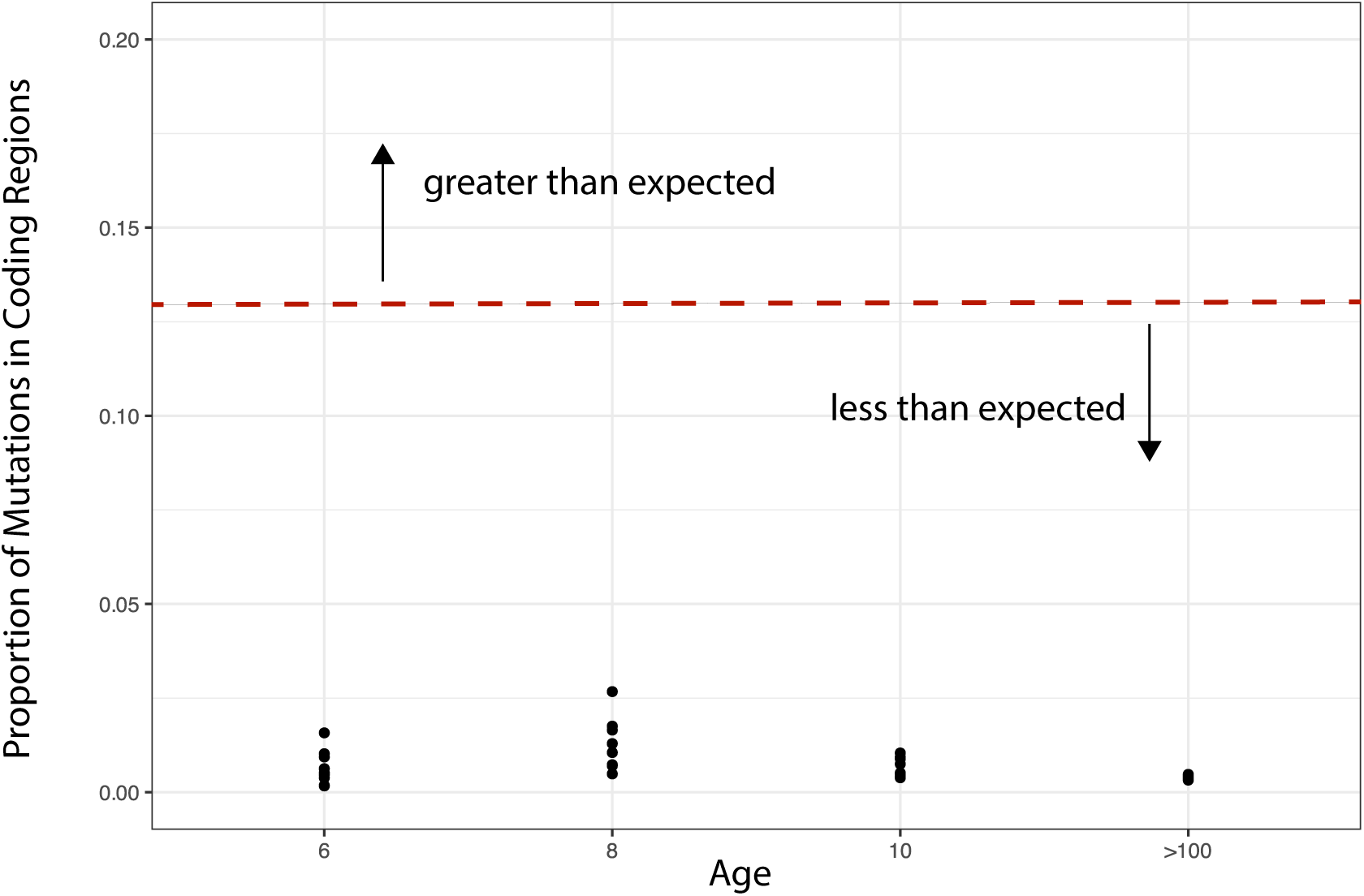
Purifying selection within a genet of *A.palmata*. Scatter plot of proportion of mutations within a module (sample) discovered in coding regions within the *A.palmata* genome across the four age classes tested in this study. Red dotted line is the proportion of the *A.palmata* genome that is coding sequences. Modules within all age classes were detected to be significantly depleted of mutations in coding regions (χ^2^, p<-0.05)

To determine whether high somatic mutation rates were associated with methylation across the coral genome, CpG islands were identified as a proxy for regions of the genome likely experiencing elevated methylation (Supplementary Data). Overall, mutations located in putative CpG islands comprised on average 0.25% of mutations identified. This was consistent across all colonies and age classes, and no significant difference was found in the proportion of mutations found in these regions with increased age (p>0.05, ANOVA) (Figure S1).

## Discussion

Here, we show that somatic genetic variation in eight *Acropora palmata* colonies from Curaçao increases with genet age. Based on the average SOGV accumulation rate of 2.798 e^-7^± 1.977e^-7^ mutations bp^-1^ module^-1^ year^-1^, we applied a somatic genetic clock, to estimate the age of a genet of uncertain age as 80.77 years, somewhat younger than previous estimates that had used a coalescent approach (228-1194 years old). In coral genets as young as six- to ten-years-old, replicate samples sequenced to a depth of at least 80x revealed high somatic genetic variation across the genomes of all genets. Although mutations were not fixed at polyp or branch levels, their allele frequencies followed a power-law distribution, similar to aging human tissues, and were not under positive selection. In fact, depletion of mutations in coding regions suggests strong purifying selection. It is likely that a hat a high-fidelity DNA repair mechanisms opposes the high substitution burden in coding regions. Surprisingly, genetic and physical distances were unrelated in samples taken along the growth axis of a large colony. These observations support the hypothesis that corals, in contrast to previous observations in vascular plants, derive new modules from a large stem cell pool, delaying fixation for a very long time. Alternatively, the observed lack of fixation and the disconnect between genetic and physical distances may be explained by multiple stem cell lineages that are shared across the colony.

### A somatic molecular clock for Acropora palmata

Previous estimates of genet age in coral populations were derived from a coalescent model of somatic mutation accumulation (Devlin-Durante et al., 2016). The method relied on a limited set microsatellite markers and could only be applied to genets with at least five living ramets. As well, microsatellites can mutate both forward and backward, generating error in the assumptions needed for a somatic genetic clock. The method is therefore skewed towards the higher range of genet ages, and it does not provide information on the population genetic processes affecting somatic mutations within colonies.

Here, we instead modify the somatic genetic clock presented in Yu et al., 2024 by focusing on mosaic SoGV. As no fixation was observed, the somatic genetic clock was instead parametrized with mosaic SoGV and calibrated based on genets of known age that can be applied to any *Acropora palmata* colony. Despite varying colony size and depth at which the colonies lived, total somatic genetic variation was consistent within the six and ten year old age classes of *Acropora palmata* colonies in Curaçao (ANOVA, p>0.05) (Figure 1b). This indicates that also mosaic SoGV feature a linear process of mutation accumulation with age, which allows us to calibrate a somatic molecular clock for *A. palmata* and apply this clock to estimate the ages of older genets of *Acropora palmata*.

Determining the precise distribution of age classes in reef populations of *Acropora palmata* is important for modeling reef futures. Natural sexual recruitment in the Caribbean has been dramatically reduced in recent years (Hughes & Tanner, 2000; Roff, 2020), calling into question the ability of these populations to adapt to changing conditions. To understand for how long this lack of recruitment has been a problem in the population, a reliable method for estimating age in younger colonies is needed and this study is a first step in that direction. Although this method is expensive and carries a high computational effort, given the increasing affordability of whole genome sequencing and availability of computational pipelines, a systematic survey should be designed to estimate the distribution of ages of remaining *Acropora palmata* in the Caribbean.

### Stem cell demography may inhibit mutation fixation within modules

Yu et al. (2024) modeled the allele frequency dynamics of somatic mutations in modular organisms and predicted that the fixation of a mutation in a modular organism such as *Acropora palmata* will be highly reliant on the demography of the stem cell pool. In the seagrass *Zostera marina*, the limited number of apical stem cells (7-12) allows for a swift and continual fixation with time as the mutations experience a strong bottleneck with the formation of a new ramet (Yu et al., 2024). However, in *A. palmata* corals, even in a colony close to a century old, no fixed mutations were detected (Figure 2). This lack of fixation of somatic mutations in old *A. palmata* genets is inconsistent with a module formation process that relies on only a few stem cells. The allele frequency spectra in *Acropora palmata* instead suggest alternative modes of module formation, alone or in combination: a large stem cell pool, multiple stem cell lines, and/or potential movement of cells across the colony that allows for the presence of the same mutation in multiple modules independent of the growth history of the modules.

First, there may be a large stem cell pool. According to the models of genetic drift within a modular organism such as a coral genet, stem cell pool (N_e_) and mutation rate are two of the most important factors influencing the allele frequencies of mutations in the population of modules (Yu et al, 2024). Similarly to populations of whole organisms, the larger the cell population size, particularly during a bottleneck event, the longer it will take to reach fixation in a part of the colony. Therefore, even across many years, the likelihood of fixation would be low in a genet of *Acropora palmata* if the stem cell pool that forms new modules is large.

Alternatively, and not mutually exclusively, stem cells either remain dormant, or somatic cells de-differentiate into stem cells, which would increase cell population size and effectively perturb any spatial correlation with the pedigree of mutation sharing. The degree of pluripotency of cells across cnidarians is highly variable. Some cnidarians, such as hydra, have three defined ‘stem’ cell lines (Holstein, 2023). Other cnidarians demonstrate pluripotency across all stem cell types that facilitate cnidarian’s impressive regeneration capabilities. The model cnidarian *Hydra* can regenerate an entire individual from a small population of cells, and the starlet sea anemone *Nematostella* can also regenerate a polyp from a section of tissue (Bossert & Thomsen, 2017; Vervoort, 2011; Vogg, Galliot, & Tsiairis, 2019). Regeneration in reef-building corals is more limited than other cnidarians but is still integral to the health and survival of a colony. When a colony is injured, or experiences partial mortality, the surrounding tissue must recruit cells to regenerate all tissue types and regrow (or ‘reskin’) the injured area (Meesters et al., 1997). Even without prior colony integration, fragments of the same genet, if placed in close enough proximity will regrow together to create one integrated colony (Forsman, Page, Toonen, & Vaughan, 2015; Highsmith, 1982). This regeneration would require the recruitment of stem-like cells to differentiate and rebuild a whole polyp and all associated tissue and cell types in the colony. Molecular signals of stem-cells were discovered at the boundaries of wounds where regeneration would occur (Levanoni, Rosner, Lapidot, Paz, & Rinkevich, 2024). Colony integration also allows corals to distribute resources across the entire colony and so repair wounds of any tissue type (Oren, Benayahu, Lubinevsky, & Loya, 2001). Colony integration may even allow for cells to be shared between modules, and this cell migration may prevent fixation of somatic mutations in subsets of modules in a genet as well as breakdown the expected relationship between genetic and physical distance along the growth axis of a genet.

### Evolutionary Consequences of Somatic Mutations in Acropora palmata

The fate of somatic mutations in animals and plants can be much more similar than previously thought. In trees, even low frequency somatic mutations are often inherited in meiotic offspring (Schmitt et al., 2024). In contrast, unitary animals often have effective separation between the soma and germline and so prevent the inheritance of somatic mutations by their meiotic offspring. In the modular animal, *Acropora palmata*, somatic mutations may not come to fixation and indeed are often maintained at lower frequency within a genet. Yet, they can still be fixed in the population of *A. palmata* genets when passing through the single-cell stage of meiotic gamete formation (Kuntz-Vasquez et al., 2022; Lopez et al., 2023; Schweinsberg et al., 2014).

The significant depletion in mutations found in coding sequences of *A. palmata* (Figure 6) might be an indicator of strong purifying selection within a genet, or of more effective repair systems acting on essential genes, or a mixture of both processes. If true, a high somatic mutation rate is controlled by a high-fidelity DNA repair mechanism. Mutation rates have been found to scale by lifespan in mammals and not body size (Cagen et al., 2022). Corals represent a unique situation where both lifespan and body size are indeterminate (Bythell et al., 2018). This allows for substantial accumulation of somatic mutations over a coral’s lifetime, thus creating a risk of mutational meltdown (Muller’s ratchet) (Antolin & Strohbeck, 1985). The maintenance of mutations in the mosaic state in *Acropora palmata* may allow the genet to generate potentially adaptive genetic diversity that can be inherited during sexual reproduction, while minimizing the mutation burden on the genet as a whole.

The patterns of somatic mutation accumulation documented here do not provide evidence that somatic mutations provide fitness advantages in *A. palmata,* but they are an important starting point to assess potential selective dynamics under different environmental conditions or in other species. Moreover, they can predict genet age in a highly clonal coral and reveal stem cell dynamics in a species for which long-term cell cultures have not successfully been established (Reyes-Bermudez & Miller, 2009). Somatic mutations create a diverse mosaic of novel alleles within a genet and in acroporid corals they can be inherited in gametes, providing additional genetic variants in species where sexual reproduction is limited to once a year. This study for the first time analyzes the shift in SoGV and the associated allele frequencies of somatic mutations with age in a reef-building coral and calibrates and applies a somatic molecular clock to determine the age of *Acropora palmata* genet.

## Acknowledgments

We would like to acknowledge Dr. Sheila Kitchen, Dr. Nicolas Locatelli, and Dr. Kathryn Stankiewicz for input on the bioinformatic analyses performed in this study.

## Data availability

Scripts and associated files are publicly available on Zenodo (https://zendodo.org/doi/10.5281/zenodo.15105057). Scripts used for somatic mutation modeling are available on github (https://github.com/jessierenton/SomaticEvolution.jl).

## Permits

All samples were collected in Curaçao under permits provided to CARMABI by the Curaçaoan Government and were imported to the United States under CITES permit 1616US784243/67 where the samples were analyzed.

## Author contributions

TLC performed sample collection, DNA extraction and sample processing, bioinformatics and writing of the paper. JR performed modeling of population dynamics within a module of a coral and wrote most of the text describing modeling methods. VC and ZD aided in sample and data collection. BW, TR, and IBB acquired funding. TLC and IBB conceived the work. TLC and IBB wrote the paper. All authors edited the paper.

## References cited

1. Ally, D., Ritland, K., & Otto, S. P. (2010). Aging in a Long-Lived Clonal Tree. PLOS Biology, 8(8), e1000454. doi:10.1371/journal.pbio.1000454

2. Antolin, M. F., & Strobeck, C. (1985). The Population Genetics of Somatic Mutation in Plants. The American Naturalist, 126(1), 52–62. doi:10.1086/284395

3. Baums, I. B., Miller, M. W., & Hellberg, M. E. (2006). GEOGRAPHIC VARIATION IN CLONAL STRUCTURE IN A REEF-BUILDING CARIBBEAN CORAL, ACROPORA PALMATA. Ecological Monographs, 76(4), 503–519. 10.1890/0012-9615(2006)076[0503:GVICSI]2.0.CO;2

4. Benjamin, D., Sato, T., Cibulskis, K., Getz, G., Stewart, C., & Lichtenstein, L. (2019). Calling Somatic SNVs and Indels with Mutect2. bioRxiv, 861054. doi:10.1101/861054

5. Bergeron, L. A., Besenbacher, S., Turner, T., Versoza, C. J., Wang, R. J., Price, A. L., … Schierup, M. H. (2022). The Mutationathon highlights the importance of reaching standardization in estimates of pedigree-based germline mutation rates. eLife, 11, e73577. doi:10.7554/eLife.73577

6. Bode, H. R. (1996). The interstitial cell lineage of hydra: a stem cell system that arose early in evolution. Journal of Cell Science, 109(6), 1155. Retrieved from http://jcs.biologists.org/content/109/6/1155.abstract

7. Bosch, T. C., Anton-Erxleben, F., Hemmrich, G., & Khalturin, K. (2010). The Hydra polyp: nothing but an active stem cell community. Dev Growth Differ, 52(1), 15–25. doi:10.1111/j.1440-169X.2009.01143.x

8. Boscolo Bielo, L., D. Trapani, M. Repetto, E. Crimini, C. Valenza, C. Belli, C. Criscitiello, A. Marra, V. Subbiah, and G. Curigliano. 2023. ’Variant allele frequency: a decision-making tool in precision oncology?’, Trends Cancer, 9: 1058–68.

9. Bossert, P., & Thomsen, G. H. (2017). Inducing complete polyp regeneration from the Aboral Physa of the Starlet Sea anemone Nematostella vectensis. JoVE (Journal of Visualized Experiments)(119), e54626.

10. Bythell, J. C., Brown, B. E., & Kirkwood, T. B. L. (2018). Do reef corals age? Biological Reviews, 93(2), 1192–1202. 10.1111/brv.12391

11. Caravagna, G., Sanguinetti, G., Graham, T. A., & Sottoriva, A. (2020). The MOBSTER R package for tumour subclonal deconvolution from bulk DNA whole-genome sequencing data. BMC Bioinformatics, 21(1), 531. doi:10.1186/s12859-020-03863-1

12. Cagan, A., Baez-Ortega, A., Brzozowska, N., Abascal, F., Coorens, T. H. H., Sanders, M. A., … Martincorena, I. (2022). Somatic mutation rates scale with lifespan across mammals. Nature, 604(7906), 517–524. doi:10.1038/s41586-022-04618-z

13. Coates, A. G., & Jackson, J. B. C. (1987). Clonal growth, algal symbiosis, and reef formation by corals. Paleobiology, 13(4), 363–378. doi:10.1017/S0094837300008988

14. Conway, J. R., Lex, A., & Gehlenborg, N. (2017). UpSetR: an R package for the visualization of intersecting sets and their properties. Bioinformatics, 33(18), 2938–2940. doi:10.1093/bioinformatics/btx364

15. Csardi, G., & Nepusz, T. (2005). The Igraph Software Package for Complex Network Research. *InterJournal*, Complex Systems, 1695.

16. Danecek, P., Bonfield, J. K., Liddle, J., Marshall, J., Ohan, V., Pollard, M. O., … Li, H. (2021). Twelve years of SAMtools and BCFtools. Gigascience, 10(2). doi:10.1093/gigascience/giab008

17. de Mendoza, A., Hatleberg, W. L., Pang, K., Leininger, S., Bogdanovic, O., Pflueger, J., … Lister, R. (2019). Convergent evolution of a vertebrate-like methylome in a marine sponge. Nature Ecology & Evolution, 3(10), 1464–1473. doi:10.1038/s41559-019-0983-2

18. Devlin-Durante, M. K., Miller, M. W., Caribbean Acropora Research, G., Precht, W. F., & Baums, I. B. (2016). How old are you? Genet age estimates in a clonal animal. Molecular Ecology, 25(22), 5628–5646. doi:10.1111/mec.13865

19. de Witte, L. C., & Stöcklin, J. (2010). Longevity of clonal plants: why it matters and how to measure it. Annals of Botany, 106(6), 859–870. doi:10.1093/aob/mcq191

20. Durante, M. K., Baums, I. B., Williams, D. E., Vohsen, S., & Kemp, D. W. (2019). What drives phenotypic divergence among coral clonemates of Acropora palmata? Molecular Ecology, 28(13), 3208–3224. doi:10.1111/mec.15140

21. Forsman, Z. H., Page, C. A., Toonen, R. J., & Vaughan, D. (2015). Growing coral larger and faster: micro-colony-fusion as a strategy for accelerating coral cover. PeerJ, 3, e1313.

22. Franco, I., & Eriksson, M. (2022). Reverting to old theories of ageing with new evidence for the role of somatic mutations. Nature Reviews Genetics, 23(11), 645–646. doi:10.1038/s41576-022-00513-5

23. Gomez-Campo, K., Sanchez, R., Martinez-Rugerio, I., Yang, X., Maher, T., Osborne, C. C., … Iglesias-Prieto, R. (2024). Phenotypic plasticity for improved light harvesting, in tandem with methylome repatterning in reef-building corals. Mol Ecol, 33(4), e17246. doi:10.1111/mec.17246

24. Highsmith, R. C. (1982). Reproduction by Fragmentation in Corals Marine Ecology Progress Series, 7, 207–226. doi:10.3354/meps007207

25. Hughes, T. P., & Tanner, J. E. (2000). RECRUITMENT FAILURE, LIFE HISTORIES, AND LONG-TERM DECLINE OF CARIBBEAN CORALS. Ecology, 81(8), 2250–2263. 10.1890/0012-9658(2000)081[2250:RFLHAL]2.0.CO;2

26. Irwin, A., Greer, L., Humston, R., Devlin-Durante, M., Cabe, P., Lescinsky, H., … Baums, I. B. (2017). Age and intraspecific diversity of resilient Acropora communities in Belize. Coral Reefs, 36(4), 1111–1120. doi:10.1007/s00338-017-1602-9

27. Kamvar, Z. N., Tabima, J. F., & Grünwald, N. J. (2014). Poppr: an R package for genetic analysis of populations with clonal, partially clonal, and/or sexual reproduction. PeerJ, 2, e281. doi:10.7717/peerj.281

28. Kitchen, S. A., Von Kuster, G., Kuntz, K. L. V., Reich, H. G., Miller, W., Griffin, S., … Baums, I. B. (2020). STAGdb: a 30K SNP genotyping array and Science Gateway for Acropora corals and their dinoflagellate symbionts. Scientific Reports, 10(1), 12488. doi:10.1038/s41598-020-69101-z

29. Klughammer, J., Romanovskaia, D., Nemc, A., Posautz, A., Seid, C. A., Schuster, L. C., … Bock, C. (2023). Comparative analysis of genome-scale, base-resolution DNA methylation profiles across 580 animal species. Nature Communications, 14(1), 232. doi:10.1038/s41467-022-34828-y

30. Moeller, M. E., Mon Père, N. V., Werner, B., & Huang, W. (2024). Measures of genetic diversification in somatic tissues at bulk and single-cell resolution. Elife, 12. doi:10.7554/eLife.89780

31. Levanoni, J., Rosner, A., Lapidot, Z., Paz, G., & Rinkevich, B. (2024). Coral Tissue Regeneration and Growth Is Associated with the Presence of Stem-like Cells. Journal of Marine Science and Engineering, 12(2), 343. Retrieved from https://www.mdpi.com/2077-1312/12/2/343

32. Li, H., & Durbin, R. (2009). Fast and accurate short read alignment with Burrows–Wheeler transform. Bioinformatics, 25(14), 1754–1760. doi:10.1093/bioinformatics/btp324

33. Lirman, D. (2000). Fragmentation in the branching coral Acropora palmata (Lamarck): growth, survivorship, and reproduction of colonies and fragments. Journal of Experimental Marine Biology and Ecology, 251(1), 41–57. 10.1016/S0022-0981(00)00205-7

34. Locatelli, N. S., Kitchen, S. A., Stankiewicz, K. H., Osborne, C. C., Dellaert, Z., Elder, H., … Baums, I. B. (2023). Genome assemblies and genetic maps highlight chromosome-scale macrosynteny in Atlantic acroporids. bioRxiv, 2023.2012.2022.573044. doi:10.1101/2023.12.22.573044

35. López-Nandam, E. H., Albright, R., Hanson, E. A., Sheets, E. A., & Palumbi, S. R. (2023). Mutations in coral soma and sperm imply lifelong stem cell renewal and cell lineage selection. Proceedings of the Royal Society B: Biological Sciences, 290(1991), 20221766. doi:doi:10.1098/rspb.2022.1766

36. Martincorena, I., Raine, K. M., Gerstung, M., Dawson, K. J., Haase, K., Van Loo, P., … Campbell, P. J. (2017). Universal Patterns of Selection in Cancer and Somatic Tissues. Cell, 171(5), 1029–1041.e1021. doi:10.1016/j.cell.2017.09.042

37. Meesters, E. H., Pauchli, W., & Bak, R. P. M. (1997). Predicting regeneration of physical damage on a reef-building coral by regeneration capacity and lesion shape. Marine Ecology Progress Series, 146, 91–99.

38. Moeller, M. E., Mon Père, N. V., Werner, B., & Huang, W. (2024). Measures of genetic diversification in somatic tissues at bulk and single-cell resolution. Elife, 12. doi:10.7554/eLife.89780

39. Oren, U., Benayahu, Y., Lubinevsky, H., & Loya, Y. (2001). COLONY INTEGRATION DURING REGENERATION IN THE STONY CORAL FAVIA FAVUS. Ecology, 82(3), 802–813. 10.1890/0012-9658(2001)082[0802:CIDRIT]2.0.CO;2

40. Paradis, E., & Schliep, K. (2019). ape 5.0: an environment for modern phylogenetics and evolutionary analyses in R. Bioinformatics, 35(3), 526–528. doi:10.1093/bioinformatics/bty633

41. Ranade, S. S., Ganea, L. S., Razzak, A. M., & García Gil, M. R. (2015). Fungal Infection Increases the Rate of Somatic Mutation in Scots Pine (Pinus sylvestris L.). J Hered, 106(4), 386–394. doi:10.1093/jhered/esv017

42. Reusch, T. B. H., Baums, I. B., & Werner, B. (2021). Evolution via somatic genetic variation in modular species. Trends in Ecology & Evolution, 36(12), 1083–1092. doi:10.1016/j.tree.2021.08.011

43. Reyes-Bermudez, A., & Miller, D. J. (2009). In vitro culture of cells derived from larvae of the staghorn coral Acropora millepora. Coral Reefs, 28(4), 859–864. doi:10.1007/s00338-009-0527-3

44. Rice, P., Longden, I., & Bleasby, A. (2000). EMBOSS: the European Molecular Biology Open Software Suite. Trends in genetics : TIG, 16(6), 276–277. doi:10.1016/s0168-9525(00)02024-2

45. Roff, G. (2020). Evolutionary History Drives Biogeographic Patterns of Coral Reef Resilience. BioScience, 71(1), 26–39. doi:10.1093/biosci/biaa145

46. Schweinsberg, M., González Pech, R. A., Tollrian, R., & Lampert, K. P. (2014). Transfer of intracolonial genetic variability through gametes in Acropora hyacinthus corals. Coral Reefs, 33(1), 77–87. doi:10.1007/s00338-013-1102-5

47. Schmitt, S., Heuret, P., Troispoux, V., Beraud, M., Cazal, J., Chancerel, É., … Tysklind, N. (2024). Low-frequency somatic mutations are heritable in tropical trees Dicorynia guianensis and Sextonia rubra. Proceedings of the National Academy of Sciences, 121(10), e2313312121. doi:doi:10.1073/pnas.2313312121

48. Schoen, D. J., & Schultz, S. T. (2019). Somatic Mutation and Evolution in Plants. Annual Review of Ecology, Evolution, and Systematics, 50(Volume 50, 2019), 49–73. 10.1146/annurev-ecolsys-110218-024955

49. Sobral, M., & Sampedro, L. (2022). Phenotypic, epigenetic, and fitness diversity within plant genotypes. Trends in Plant Science, 27(9), 843–846. 10.1016/j.tplants.2022.06.008

50. Tomiak, P. J., Andersen, M. B., Hendy, E. J., Potter, E. K., Johnson, K. G., & Penkman, K. E. H. (2016). The role of skeletal micro-architecture in diagenesis and dating of Acropora palmata. Geochimica et Cosmochimica Acta, 183, 153–175. 10.1016/j.gca.2016.03.030

51. Van Oppen, M. J. H., Souter, P., Howells, E. J., Heyward, A., & Berkelmans, R. (2011). Novel Genetic Diversity Through Somatic Mutations: Fuel for Adaptation of Reef Corals? Diversity, 3(3), 405–423. doi:10.3390/d3030405

52. Vasquez Kuntz, K. L., Kitchen, S. A., Conn, T. L., Vohsen, S. A., Chan, A. N., Vermeij, M. J. A., … Baums, I. B. (2022). Inheritance of somatic mutations by animal offspring. Science Advances, 8(35), eabn0707. doi:doi:10.1126/sciadv.abn0707

53. Vervoort, M. (2011). Regeneration and development in animals. Biological Theory, 6, 25–35.

54. Vogg, M. C., Galliot, B., & Tsiairis, C. D. (2019). Model systems for regeneration: Hydra. Development, 146(21), dev177212.

55. Williams, M. J., Werner, B., Barnes, C. P., Graham, T. A., & Sottoriva, A. (2016). Identification of neutral tumor evolution across cancer types. Nature Genetics, 48(3), 238–244. doi:10.1038/ng.3489

56. Xia, J., Han, L., & Zhao, Z. (2012). Investigating the relationship of DNA methylation with mutation rate and allele frequency in the human genome. BMC Genomics, 13(8), S7. doi:10.1186/1471-2164-13-S8-S7

57. L., Boström, C., Franzenburg, S., Bayer, T., Dagan, T., & Reusch, T. B. H. (2020). Somatic genetic drift and multilevel selection in a clonal seagrass. Nature Ecology & Evolution, 4(7), 952–962. doi:10.1038/s41559-020-1196-4

58. Yu, L., J. Renton, A. Burian, M. Khachaturyan, T. Bayer, J. Kotta, J. J. Stachowicz, K. DuBois, I. B. Baums, B. Werner and T. B. H. Reusch (2024). “A somatic genetic clock for clonal species.” Nature Ecology & Evolution 8(7): 1327–1336.

