## Supplementary Data for "Mosaic accumulation of somatic genetic variation and estimates of age in the long-lived reef-building coral *Acropora palmata*"

*Identification of CpG Islands*

CpG rich regions of the *Acropora palmata* genome were identified using the cpgreport function in the program EMBOSS (Rice, Longden, & Bleasby, 2000). A score threshold of 28 was used. There is some evidence that somatic mutations are associated with highly methylated regions of the genome. To determine if there was a statistically significant association between the identified mutations and CpG regions, first, regions with elevated CpG content were identified. Then, identified mutations were mapped to regions of CpG content. The proportions of mutations identified in CpG islands vs. proportions of mutations identified across the rest of the genome were compared statistically using R v.4.3.2 (Figure S1).

*Modeling Population Expectations*

To estimate what the most important parameters were in the accumulation and fixation of mutations in *Acropora palmata*, we implement a stochastic model of a clonal organism as follows. The organism consists of a collection of modules formed of (stem) cells. In the case of coral, the module would represent a single polyp. In our simulations we follow a single module that grows from a single cell by cell division rate at rate $b$ until it reaches size $N$ and then enters a homeostatic phase in which cells can divide asymmetrically (resulting in a single child cell) or symmetrically (producing two child cells, a random stem cell is then removed). These homeostatic divisions also occur at a rate $b$. Mutations arise at cell division (during growth and homeostasis), the number of which is Poisson distributed with mean $u$, and every mutation is assumed to be unique (infinite sites model). New modules are formed at a rate $r$, by module splitting (the module splits into two modules of size $N_0$ and $N - N_0$) or module branching ($N0$ cells are sampled without replacement and copied to form a new module, leaving the original module unchanged). In these simulations we have considered a population containing a single module. Thus, after a new module is formed, we discard either the parent or child module at random. This is the same model implemented in (Yu et al., 2024). We perform simulations with neutral selection for a fixed time and then compute the variant allele frequency (VAF) spectra. We are interested in how different parameters affect the VAF spectra, and particularly whether they conform to power law regimes that have been established in single level population models. In a single-level population exhibiting exponential growth the VAF spectrum should follow a $1/f^2$ law. Conversely, in a constant size single-level population model, such as a Moran process, the VAF spectrum should have a 1/f form. However, while the $1/f^2$ law is adhered to immediately in the case of exponential growth, the 1/f law for constant populations is only achieved in the long-time limit. Thus, if a population grows exponentially and then switches to a constant population size, it will take time to move from $1/f^{2 \text{ to}}$ 1/f, and this will initially happen in the lower part of the VAF spectrum. We consider whether our two-level modular population conforms to these laws, and how various parameters affect the results. We have set cell division rate $b$ = 120/year and mutation rate $u$ =0.01. Other parameters are varied, as is the choice between symmetric or asymmetric cell division.

The VAF spectra for asymmetric cell division was modeled with slow (*r*=5 year) and fast (*r=80/year)* formation, respectively (Figure S3, Figure S4). In both cases, when the bottleneck at module formation is small (e.g. $N_0$ =1, $N_0$ =10) the VAF spectrum clearly follows a $1/f^2$ distribution, indicating that the dominant mode of module dynamics is exponential growth. For intermediate bottleneck size ($N_0$ =100) this $1/f^2$ trend is still apparent, particularly in the lower part of the VAF spectrum. However, the VAF spectrum begins to curve up slightly at higher frequencies. For $N_0$ = 1000 only extremely low allele frequencies fall on the $1/f^2$ line, and the line quickly curves upwards to follow a 1/f trend, indicative of a constant population. For $N_0$ = 5000 the curve follows a 1/f, after a sufficiently long time has elapsed. In younger modules, e.g. 5 years, the lines still curve down towards the $1/f^2$ line, as is observed in constant populations under a Moran process. This is because it takes time to reach the equilibrium 1/f distribution in a constant population.

Essentially then populations with a large bottleneck mimic the constant population model such as a Moran process, while in populations with a small bottleneck, the exponential growth signatures dominate. A transitional regime is apparent for intermediate bottleneck sizes.

The VAF spectra for was also modeled for symmetric cell division with slow (*r = 5/year)* and fast (*r=80/year)* module formation rate, respectively. In the former case, cells undergo many symmetric divisions before module branching occurs. Thus, the ongoing Moran process that occurs within modules dominates, and even for small bottlenecks the VAF spectra follows a 1/f distribution (Figure S5, Figure S6).

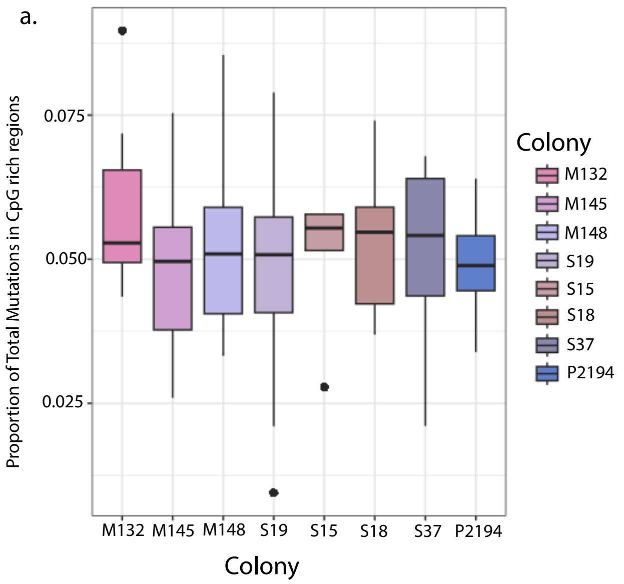


**Figure S1*: *Potential bias introduced through the mutation of methylated gene bodies. (A)** Distribution of identified somatic mutations among CpG enriched regions within each genet analyzed in this study. CpG islands were identified using the program EMBOSS. **B)** Distribution of the proportion of identified somatic mutations among age classes. There was no significant difference in the proportion of mutations identified in CpG islands across age classes

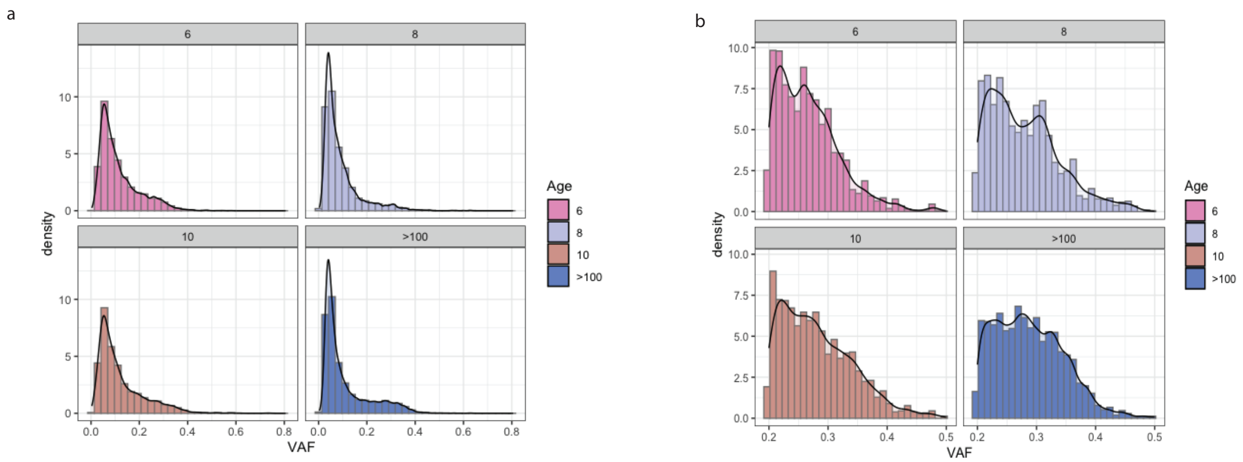


**Figure S2: Variant allele frequency distributions across age classes.** (a) Distribution of variant allele frequencies across all age classes of *Acropora palmata* tested in this study. (b) Distribution of variant allele frequencies within the mosaic range (0.2-0.5).

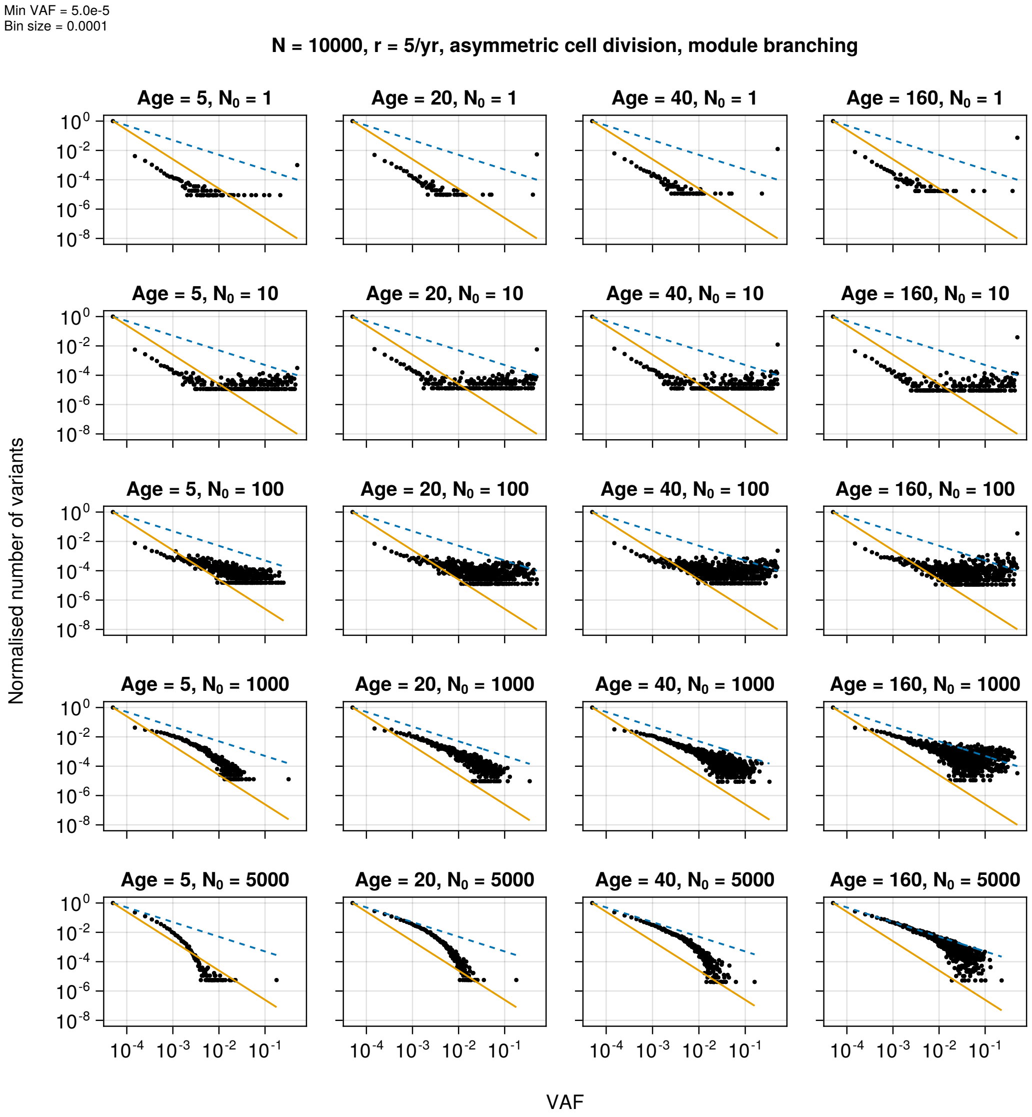


**Figure S3:** VAF spectra for a single module in a simulated clonal organism with asymmetric cell division and r=5/yr. Simulation data (black dots) are normalized so that the first data point has 'normalized number of variants' equal to one. Solid yellow line is $1/f^2$ and blue dashed line is $1/f$ (shifted to pass through the first simulation point). Data for each panel is obtained from 20 simulation runs.

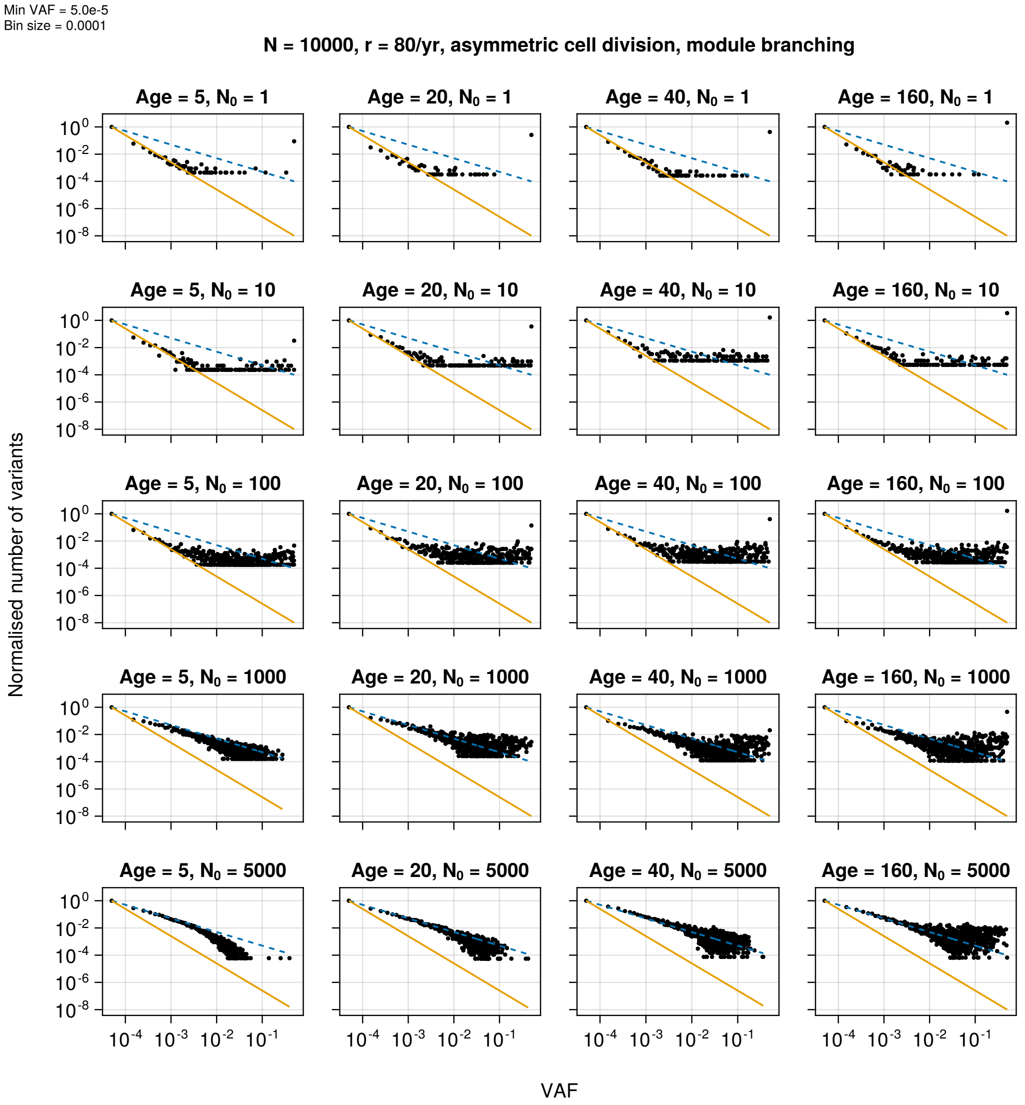


**Figure S4:** VAF spectra for a single module in a simulated clonal organism with asymmetric cell division and r=80/yr. Simulation data (black dots) are normalized so that the first data point has 'normalized number of variants' equal to one. Solid yellow line is 1/f2 and blue dashed line is 1/f (shifted to pass through the first simulation point). Data for each panel is obtained from 20 simulation runs.

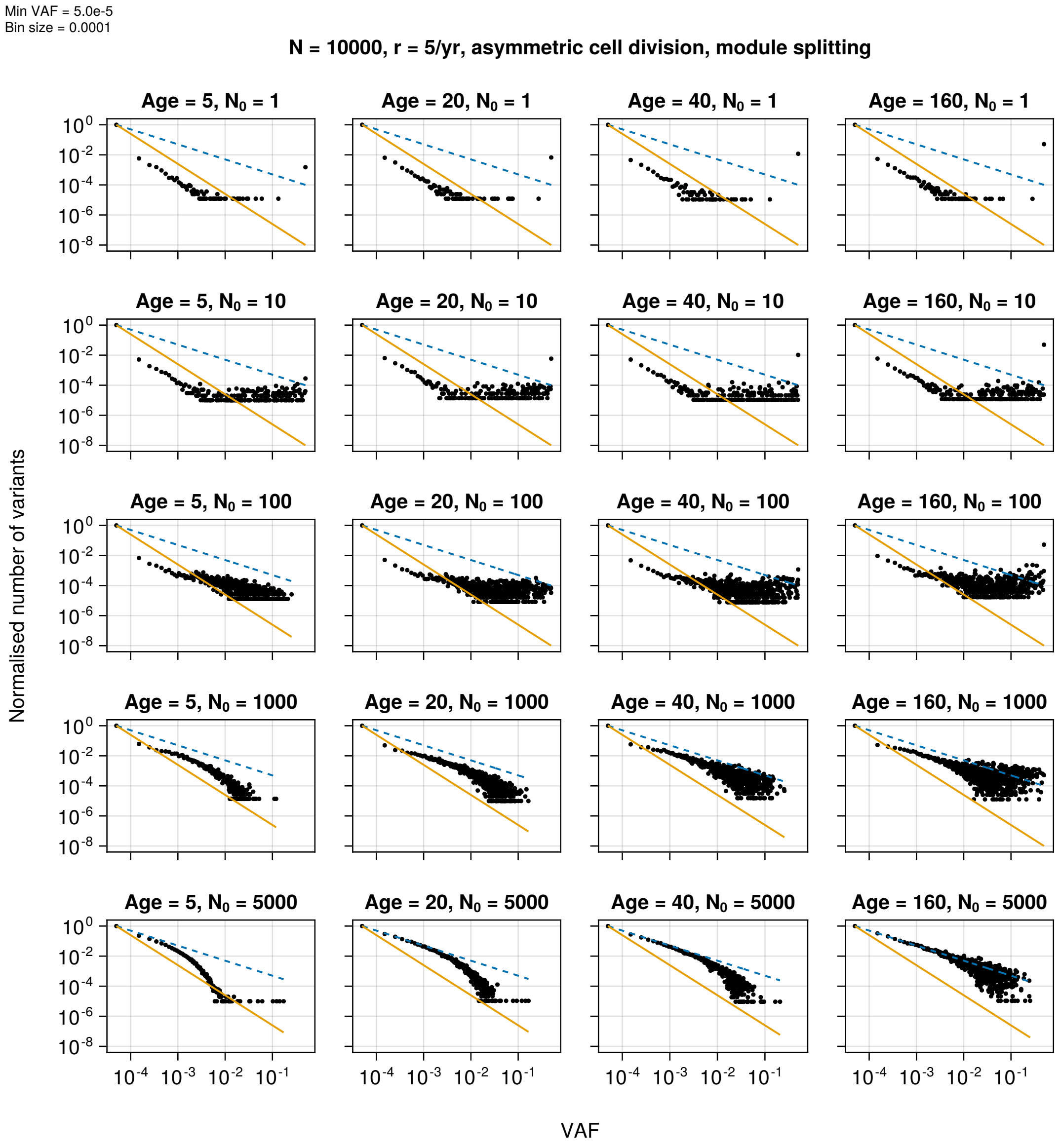


**Figure S5:** VAF spectra for a single module in a simulated clonal organism with symmetric cell division and r=5/yr. Simulation data (black dots) are normalized so that the first data point has 'normalized number of variants' equal to one. Solid yellow line is $1/f^2$ and blue dashed line is 1/f (shifted to pass through the first simulation point). Data for each panel is obtained from 20 simulation runs.

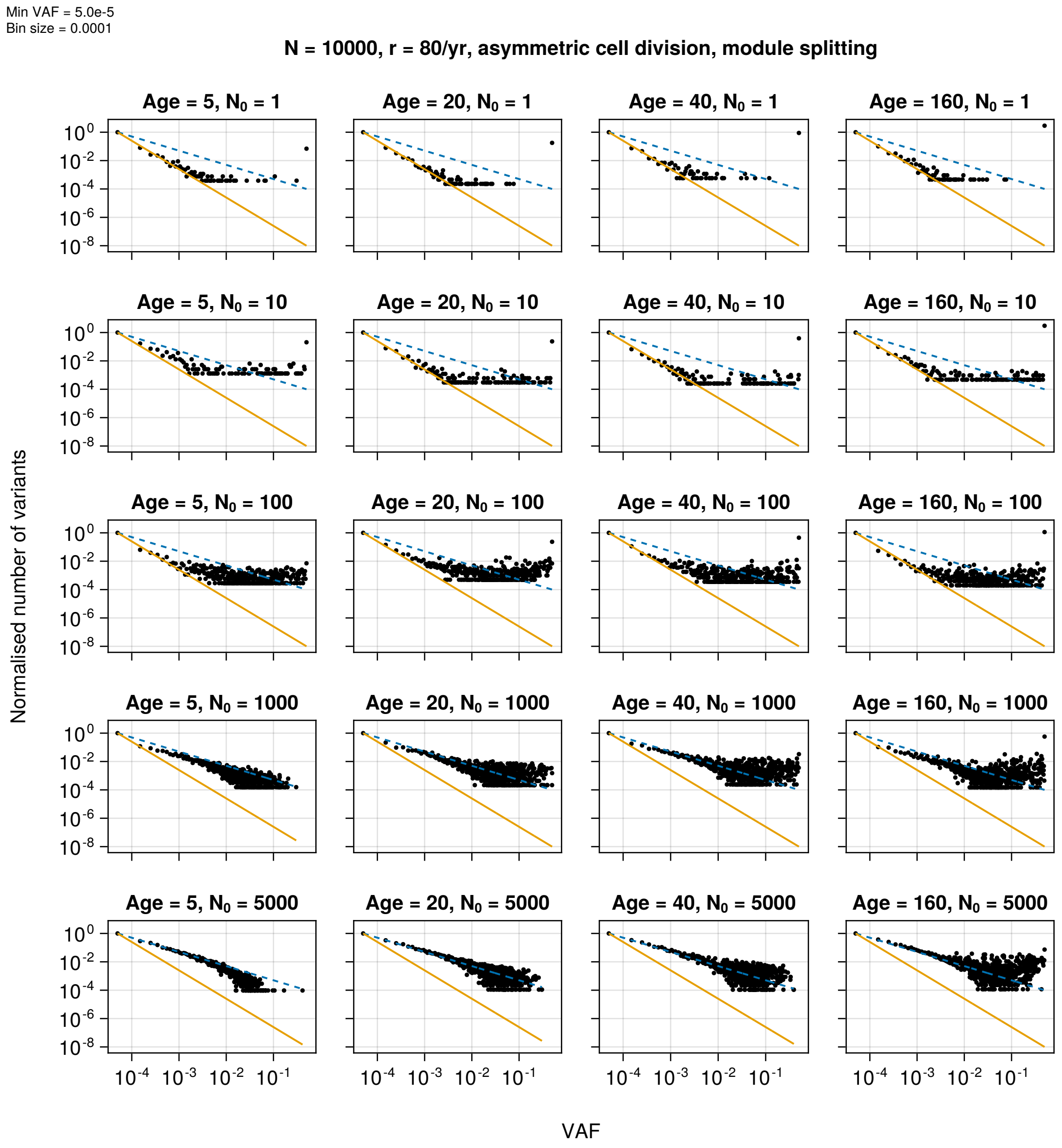


**Figure S6:** VAF spectra for a single module in a simulated clonal organism with asymmetric cell division and r=80/yr. Simulation data (black dots) are normalized so that the first data point has ‘normalized number of variants’ equal to one. Solid yellow line is $1/f^2$ and blue dashed line is $1/f$ (shifted to pass through the first simulation point). Data for each panel is obtained from 20 simulation runs.

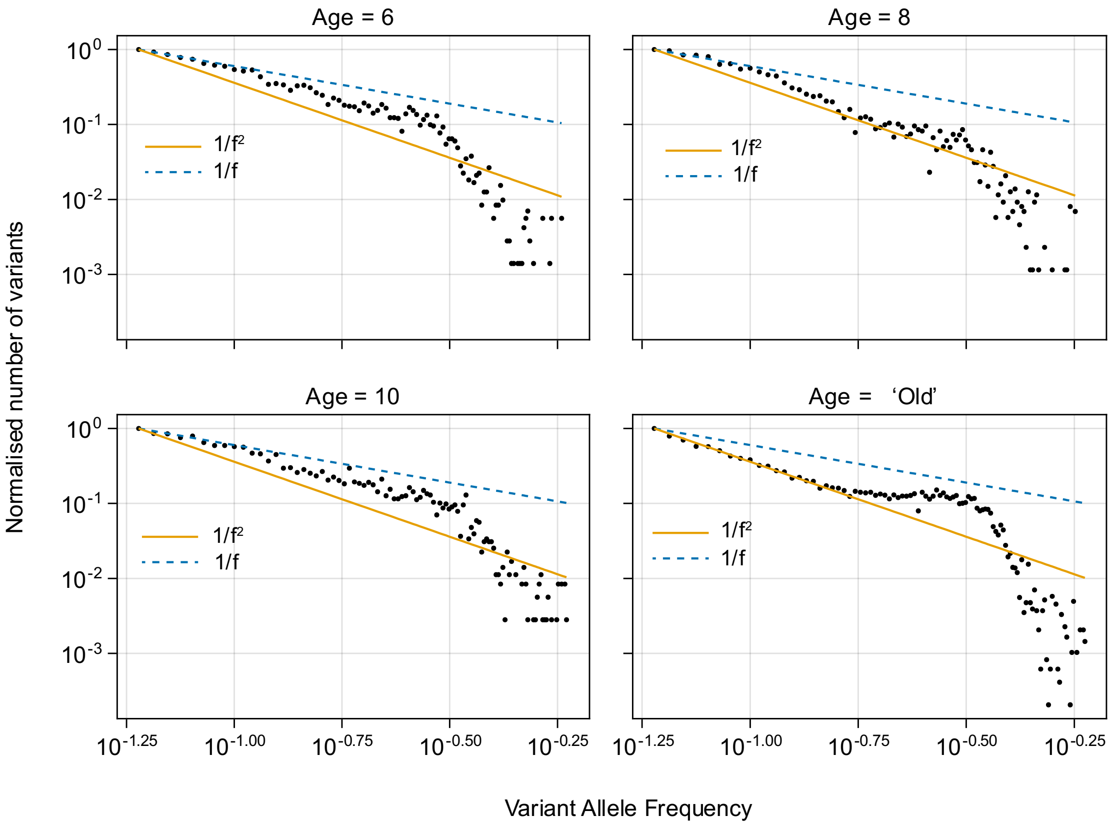


**Figure S7: Distribution of empirical variant allele frequencies across four age classes as compared to the theoretical neutral power law distributions.** X-axes represent the Variant allele frequency (VAF), with increasing VAF towards the right side of the graph. Graphs are presented for each age class (6, 8, 10 years old and an old colony of unknown age. Theoretical 1/f (blue dotted line) and $1/f^2$ distributions (yellow solid line) are represented in each figure for comparison. All age classes (6, 8, 10) except for the colony of unknown age followed a 1/f distribution at the lower variant allele frequencies.
